# Unpredictable chronic stress produces a mixed depression/anxiety behavioral profile in zebrafish

**DOI:** 10.64898/2026.09.10.750503

**Authors:** Thaeny Regina Ribeiro Rodrigues, Aurora Rúbria Batista Pantoja, Adriano Oliveira Silva de Meneses, João Alphonse Heymbeeck, Carolainy Lorrany Duarte de Sousa, Monica Lima-Maximino, Caio Maximino

## Abstract

Stress is a physiological, adaptive, and preliminary response of the organism to situations of threats and instability caused by environmental and psychological stressors. The chronicity and predictability of stressors are important variables in determining neurobehavioral outcomes. While unpredictable chronic stress (UCS) protocols typically induce depressive-like phenotypes in rodents, in zebrafish the effects of these protocols are assessed using anxiety-like outcomes. This study aimed to analyze the effects of an unpredictable UCS on depressive-type behavioral parameters and brain lipid peroxidation in zebrafish. A 14-day UCS protocol was used, with 10 different types of stressors administered twice daily in a random manner, using draws, for greater unpredictability. Animals underwent the Novel Tank Test (anxiety-like outcome) and the Social Preference Test (depressive-like outcome) 24 hours after the end of the UCS protocol. Biochemical data were collected using a thiobarbituric acid reactive substances (TBARS) assay to quantify malondialdehyde (MDA), an indicator of lipid peroxidation. The findings suggest that prolonged exposure to stressors causes damage to the organism, with increased anxiety and decreased social preference, highlighting zebrafish as an efficient model in the investigation of the neurobiological and neuropathological mechanisms of stress.

---

The etymology of the word *stress* derives from the Latin term *stringere* (to squeeze, compress, or bind), which was originally used in the field of physics. However, the concept was introduced into biology by the Canadian physiologist Hans Selye (1936, 1950), who characterized stress as a set of biochemical and physiological responses of the organism to any demand, whether caused by or resulting from favorable and/or unfavorable conditions. This set of reactions was termed the General Adaptation Syndrome (GAS), which comprises nonspecific responses triggered by the organism when exposed to stimuli that threaten homeostasis (Viner, 1999).

With the advancement of scientific research on the consequences of stressors on organisms, studies began to investigate the development of diseases resulting from continuous exposure to stress. It has been observed that biochemical and behavioral responses associated with stress promote adaptive reactions. However, when the threshold of adaptation is exceeded, deleterious effects on the organism may emerge, resulting in a state of allostatic load and increasing susceptibility to the development of diseases (Herman, 2013). Continuous exposure to chronic stressors may lead to maladaptive functioning of the hypothalamic-pituitary-adrenal (HPA) axis, resulting in neuroendocrine dysfunctions, particularly in cortisol regulation, the main glucocorticoid hormone associated with the stress response. Such dysfunction may increase the risk of developing pathological conditions, including depressive disorders (Sharan & Vellapandian, 2024).

Depression, as a disorder associated with psychological and behavioral factors, is linked to physiological alterations in the HPA axis and, consequently, to changes in cortisol concentrations in plasma, urine, and cerebrospinal fluid (CSF), although this phenomenon is not yet fully understood (Juruena et al., 2018; Schumacher & Santambrogio, 2023). Studies indicate that chronic stress, particularly when unpredictable and uncontrollable, may alter the neurobiology of the brain. Structures such as the hippocampus and amygdala, which are directly involved in emotional regulation and the neurochemical processing of stress, are affected (McEwen, 2007). The unpredictability of the stressor intensifies its detrimental effects on the organism, as the individual has no precise control over the occurrence of a new stressful episode, which prevents effective adaptation and maintains a continuous state of alertness. The interaction among emotional, cognitive, and neurobiological factors may contribute to the emergence of depressive symptoms, such as loss of interest, feelings of hopelessness, extreme fatigue, and anhedonia (Thoits, 2010).

Accordingly, the use of animal models to investigate the effects of stress on organisms has increased, particularly because they offer a favorable cost-benefit ratio for behavioral and neurophysiological studies. An innovative and valuable model that has attracted increasing attention from the scientific community is the zebrafish (*Danio rerio*), which has been widely used in neuroscience research, particularly for understanding stress-related conditions and the development of mental disorders (Yang et al., 2025). When subjected to an unpredictable chronic stress (UCS) protocol, zebrafish exhibit behavioral, biochemical, and molecular alterations that may be associated with conditions resulting from chronic exposure to stressors. This small freshwater fish possesses characteristics that make it an advantageous model for biomedical research, providing important insights into the mechanisms underlying stress and affective disorders (Gallas-Lopes et al., 2023; Yang et al., 2025).

Among the biochemical alterations associated with stress, lipid peroxidation is particularly relevant. This process results from the interaction of reactive oxygen species (ROS) with lipids in cell membranes, particularly polyunsaturated fatty acids. Lipid oxidation compromises cellular integrity and represents an important marker of oxidative stress. In zebrafish exposed to CUS, a significant increase in end products of lipid peroxidation, such as malondialdehyde (MDA), has been observed, indicating oxidative cellular damage, which may have detrimental effects on the central nervous system (Marcon et al., 2018; Mocelin et al., 2019; Pinto et al., 2026). Treatment with antioxidants have demonstrated a causal relationship between CUS and physiological and behavioral changes, suggesting an interaction between cellular metabolism and the development of disease (Marcon et al., 2019; Mocelin et al., 2019). These findings support the hypothesis that metabolic dysfunction, particularly an increase in reactive oxygen species (ROS), may act as a central biological mechanism in the onset and persistence of pathological conditions (Marcon et al., 2018, 2019; Mocelin et al., 2019).

Nonetheless, the literature overwhelmingly reports anxiety-like behavior as the primary behavioral outcome of unpredictable chronic stress (UCS) in zebrafish, with elevated cortisol as a consistent physiological correlate (Gallas-Lopes et al., 2023). A systematic review and meta-analysis of 38 studies found that UCS reliably increases anxiety/fear-related behavior and cortisol while decreasing locomotor function, with no significant summary effect for social behavior (Gallas-Lopes et al., 2023). Multiple independent groups confirm this anxiogenic profile using the novel tank diving test, where stressed fish show reduced vertical exploration, increased bottom-dwelling, and prolonged freezing (Chakravarty et al., 2013; Golla et al., 2020; Neves et al., 2026; Piato et al., 2011; Wu et al., 2023). One key study directly compared stress durations and found that 5-week UCS induces anxiety-like behavior in the novel tank test (NTT) with elevated cortisol and pro-inflammatory markers, whereas 12-week UCS induces depression-like behavior in the tail immobilization test with lowered cortisol, impaired serotonin turnover, and inflammatory activation (Kotova et al., 2025). The zebrafish tail immobilization test is based on the rodent tail suspension test; however, the interpretation of escape/immobility behaviors in the latter test as indicative of antidepressant activity has been criticized on the basis of a lack of face, construct, and predictive validity (Carvalho et al., 2021; Trunnell et al., 2024). Additional studies frame UCS as modeling depression by demonstrating neurochemical parallels (reduced whole-brain serotonin, 5-HIAA, and dopamine) and by reporting that UCS produces behavioral and molecular changes similar to those in depressed patients (Fulcher et al., 2017; Kotova et al., 2025; Marcon et al., 2016). The original UCS protocol paper itself noted impaired learning alongside anxiety, broadening the phenotype beyond pure anxiety (Piato et al., 2011). However, the lack of behavioral endpoints that explicitly model depressive-like phenotypes, such as anhedonia, represents an important limitation in the use of UCS protocols in zebrafish.

One potential candidate for an anhedonia-like phenotype in zebrafish is social preference. Differently from laboratory rodents, which usually show social withdrawal when exposed to novel conspecifics (File & Hyde, 1978), zebrafish usually present social approach responses (Barba-Escobedo & Gould, 2012; Ogi et al., 2021). Moreover, the sight of conspecifics can be used as a reinforcer in associative learning in zebrafish (Al-Imari & Gerlai, 2008), and acute social isolation increases preference and decreases dopaminergic activity (Shams et al., 2017). Moreover, blocking dopamine D1 receptors disrupts social preference in zebrafish, further implicating the dopaminergic system in this function (Scerbina et al., 2012). While variations in the social preference test exist (Ogi et al., 2021), a two-stage protocol in which both the absence/presence of conspecifics and the novelty of the stimulus are manipulated helps to discriminate between general social motivation/affiliation and social recognition/social novelty preference. In this version, the first stage (social investigation) measures whether the zebrafish prefers any conspecific over an empty tank, reflecting basic social motivation/sociability, while the second stage (social novelty) measures whether the zebrafish prefers a novel conspecific over the now-familiar conspecific from Stage 1, requiring the fish to recognize the familiar individual and exhibit a preference for social novelty. Thus, the social investigation/social novelty version of the test allows to discriminate between social withdrawal/reduced social motivation (a putative index of anhedonia) and social recognition memory and novelty-seeking. If a given manipulation, such as UCS, is inducing an anhedonia-like state, social approach should decrease during social investigation, while still being able to discriminate between conspecifics during social novelty. Nonetheless, in a one-stage version of the test, which provided a choice between a shoal (10 fish) and an empty tank, UCS did not change social preference (Neves et al., 2026), putting into question whether social preference can be used as an anhedonia-like endpoint in zebrafish. In the present study, we report the effects of UCS on anxiety-like behavior in the NTT, replicating previous experiments, and also show that the same protocol decreases social approach during social investigation (SI), increases preference for the known conspecific during social novelty (SN), and decreases preference for the unknown conspecific during SN. Moreover, we show that UCS elicits brain lipid peroxidation. We argue that this is consistent with a mixed depression/anxiety phenotype.

## 2. Materials and methods

### 2.1. Animals and housing

Forty zebrafish (*Danio rerio*) were used in the experiment. Considering an effect size of *d* = 0.79 (half the size reported by the meta-analysis made by Gallas-Lopes et al. (2023) for anxiety-like behavior in a 14-day protocol), 80% power, and α = 0.05, a model with two groups results in a minimum sample size of 19 animals/group. Animals were sexed after the end of the experiments by examination of the gonads (Siegfried & Steinfeld, 2021); the final male:female ratio was 6:31. No significant differences were observed *a posteriori* in the distribution of sexes across treatments (χ²_[df= 1]_ = 2.998, p = 0.168). Other baseline characteristics of the subjects (total length, standard length, body weight, interorbital distance, head length, mouth length, maximum body depth, anal fin base, and dorsal fin base) were recorded before experiments began; sex and morphometric measurements are available in Supplementary File S1. All animals were drug- and test-naive. The animals were maintained in collective tanks for two months prior to the experiment under a 12 h light/12 h dark photoperiod. Regarding water quality parameters, the temperature was maintained at 25 ± 3 °C, and the pH ranged from 7.5 to 8.0. The water was chlorine-free and continuously oxygenated and filtered using pumps. Nitrite levels ranged from <0.01 to 0.50 ppm, while ammonia levels ranged from 0.25 to 1.00 ppm.

### 2.2. Unpredictable chronic stress (UCS) protocol

The unpredictable chronic stress (UCS) used in this study is an adaptation of the protocol from the studies by Piato et al. (2011), Manuel et al. (2014) and Lima-Maximino et al. (2025). The protocol uses unpredictable and uncontrollable stressors for a period of 14 days, with the animal being exposed to a stressor twice a day (Table 1). Before beginning experiments, animals were randomly allocated to each group (control vs. UCS) with the help of an online tool (https://www.randomizer.org/); the sequence of allocation was randomized using the same tool. To avoid batch or tank effects (Stewart et al., 2015), animals for all groups were drawn at random from 4 different tanks located in the same facility; thus, randomization was stratified based on tank of origin. While experimenters that applied the protocol were unblinded, caretakers were blind to treatment during the experiment, and data analysts were blinded by resorting to coded videos.

**Table 1.** Protocol for unpredictable chronic stress in zebrafish.

| Day of the week | Week 1 | Week 2 |
| --- | --- | --- |
| 1 | 09:50 - Alarm substance<br>15:10 - Overcrowding | 11:00 - Heating<br>16:00 - Alarm substance |
| 2 | 08:40 - Low water level<br>14:20 - Fasting | 08:30 - Chasing<br>13:00 - Tank change |
| 3 | 10:15 - Exposure to predator animation<br>16:00 - Heating | 10:00 - Overcrowding<br>15:00 - Cooling |
| 4 | 08:30 - Overcrowding<br>13:00 - Chasing | 10:40 - Restraint<br>14:45 - Tank change |
| 5 | 10:30 - Heating<br>15:30 - Cooling | 09:30 - Chasing<br>15:30 - Cooling |
| 6 | 09:00 - Low water level<br>14:00 - Tank change | 09:00 - Crowding<br>14:00 - Low water level |
| 7 | 10:00 - Cooling<br>15:10 - Restraint | 08:00 Exposure to predator animation |
|  |  | 13:40 - Fasting |

### 2.3. Behavioral Tests

#### 2.3.1. Novel Tank Test (NTT)

The novel tank test (NTT) evaluates zebrafish responses to a potential threat in a novel environment; in the NTT, animals introduced to a novel tank first present a diving response (i.e., swim to the bottom) and gradually habituate during a 6-min. session (Bencan et al., 2009; Levin et al., 2007; Wong et al., 2010). The main outcome of the test is geotaxis (expressed either as time on bottom of the tank or time on top of the tank), a variable that is sensitive to drug treatments (Kysil et al., 2017); importantly, it is also sensitive to UCS (Gallas-Lopes et al., 2023).

The protocol used in this study was an adaptation of the protocol by Barra et al. (2023). Briefly, the animals were transferred individually to a transparent glass tank measuring 21 cm x 26 cm x 16 cm (height x width x length), with three of its four sides covered with white paper (only the front side was left uncovered), in order to prevent the animal from receiving interference from the external environment; the base of the tank was covered with a white Styrofoam sheet, thus decreasing the possible impacts of vibration as well as providing a white bottom color. The test started when animals were transferred to the tank, allowing them to engage in exploratory behavior for a period of 6 minutes. This activity was filmed using Moto G 14 and Redmi Note 13 Pro cellphones (50 MP; 480p resolution at 30 fps), positioned in front of the tank. The environmental conditions of light and sound maintained during the experiment averaged 443 ± 44 and 64 ± 4 dB, respectively, with constant Gaussian white noise present throughout the tests.

Subsequently, during the video analysis phase, conducted using TheRealFishTracker software (v 0.4.0, https://www.dgp.toronto.edu/~mccrae/projects/FishTracker/), the following endpoints were considered: Time spent by the fish at the bottom third of the tank (main endpoint); Time spent by the fish at the top third of the aquarium; Erratic swimming, extracted as absolute turn angle; and Average swimming speed of the fish. Differently from other protocols, here “bottom” and “top” are considered as one third of the tank, and not half; this has the effect of enhancing the assay’s sensitivity by preventing ceiling effects and providing a wider dynamic range to distinguish severe anxiety from moderate stress. Since video transcription was automated, the need for blinding to reduce outcome evaluation bias is reduced; nonetheless, the data analysts who processed the videos were blinded to treatment by using codes in file names and identification. Codes were unblinded only after the end of data analysis.

#### 2.3.2. Social Preference Test (SPT)

The social preference test was divided in two stages, social interaction (SI) and social novelty (SN) tests, and carried out based on the protocol described by Barba-Escobedo and Gould (2012). These stages aim to analyze the tendency of zebrafish to approach co-specifics and investigate novel social situations. 74 adult zebrafish were used as conspecific stimuli. These were drawn from different tanks than the hometank of the focal fish, and therefore focal fish had no prior contact with them. Immediately after the end of the NTT, the focal fish was transferred to the test tank where it remained for 3 min for acclimatization. For both SI and SN, three tanks were used: the main tank (15 cm x 25 cm x 20 cm; width, length, height), for the focal animal, and two other tanks for the conspecifics (15 cm x 17 cm x 16 cm; width, length, height). Both sides of the main tank were separated from the lateral tanks with barriers to prevent visual access. Once the fish’s acclimatization period was over, the barriers were removed allowing the focal fish visual access to the conspecifics. In SI, the behavior of 1 focal animal was evaluated for an initial period of 6 min, during which it had visual access to an unknown conspecific and an empty tank. After that time interval, barriers were lowered, a new conspecific was transferred to the empty tank, and the barriers were again removed to allow the focal fish visual access to the conspecifics. The behavior was observed again for 6 minutes, consisting in the social novelty test (SN). The conspecific that was present in the former stage was labeled as “old stranger”, while the conspecific that was introduced in the empty tank was labeled as “new stranger”. In both stages, the conspecific stimulus was a female zebrafish, confirmed by body morphology and, later, gonadal histology; female zebrafish stimuli are reported to produce stronger preferences than male stimuli (Ogi et al., 2021). In both the social interaction and social novelty stages, behavior was filmed and later analyzed using TheRealFishTracker software (v. 0.4.0; https://www.dgp.toronto.edu/~mccrae/projects/FishTracker/). The following variables were analyzed: Time near S1 (conspecific during SI, “old stranger” during SN), in s; Time near S2 (empty tank during SN, “new stranger” during SN), in s; Erratic swimming, extracted as absolute turn angle; Geotaxis, extracted as maximum distance to the bottom of the tank, in cm; and Swimming speed, in cm/s.

### 2.4. Lipid peroxidation assay

Brain and head kidney lipid peroxidation rates were analyzed using the TBARS (thiobarbituric acid-reactive substances) assay, which allows for the quantification of malondialdehyde (MDA), a byproduct generated by lipid oxidation in the samples. The protocol was adapted for zebrafish based on the methods described by Sachett et al. (2020). The concentrations of TBARS-reactive substances were adjusted according to the amount of protein present in each sample, as determined by the Bradford method (Zor & Selinger, 1996).

### 2.5. Statistical Analysis

Before analyses, data were inspected for violations in normality (Shapiro-Wilk’s test) and homocedasticity (Brown-Forsythe’s test); when assumptions were not met, analyses were made using nonparametric tests. For the NTT and lipid peroxidation assays, data were analyzed using Student’s t-tests for independent samples or Mann-Whitney’s tests. For the SPT, data were analyzed using repeated measures analyses of variance (ANOVA), with state (SI vs. SN) as within-subjects factor and stressor (control vs. UCS) as between-subjects factor; when effects were statistically significant, post-hoc Holm’s tests were applied. A significance level of p < 0.05 was used in all analyses performed. Effect sizes for *t*-tests and post-hoc tests were reported as Cohen’s *d*; effect sizes for Mann-Whitney tests were reported as biserial rank correlations; effect sizes for ANOVAs were reported as partial η² (η²p). A survival curve was also drawn based on the time to a lethal event in each group; these were analyzed using Mantel-Haenszel tests. To assess correlations between behavioral and biochemical variables, Pearson’s *r* was applied to the data separated by group. While sex was not a variable of interest in the present study, it was included in complementary statistical analysis (Supplementary File S2). All analyses were made using JASP (JASP Team, 2025).

## 3. Results

Three deaths occurred in the UCS group, and none in the control group. All deaths in the UCS group occurred between days 4 and 5 of the stress protocol. No significant differences between groups were found in survival curves (χ²_[df= 1]_ = 3.005, p = 0.083).

In the novel tank test, statistically significant differences were observed between the groups for the variables time in the bottom (t_[df= 35]_ = -4.748, p < 0.001, *d* = -1.566; Figure 1A) and time in the top (U = 287.5, p < 0.001, *d* = -0.6912; Figure 1B). These findings suggest that animals exposed to the UCS protocol were more likely to spend more time at the bottom of the apparatus and less time at the top compared with CTRL animals. The other variables did not show statistically significant differences between the groups, including erratic swimming (t_[df= 35]_ = 0.837, p = 0.408, *d* = 0.276; Figure 1C) and swimming speed (t_[df= 35]_ = 0.465, p = 0.645, *d* = 0.153; Figure 1D).

**Figure 1.**
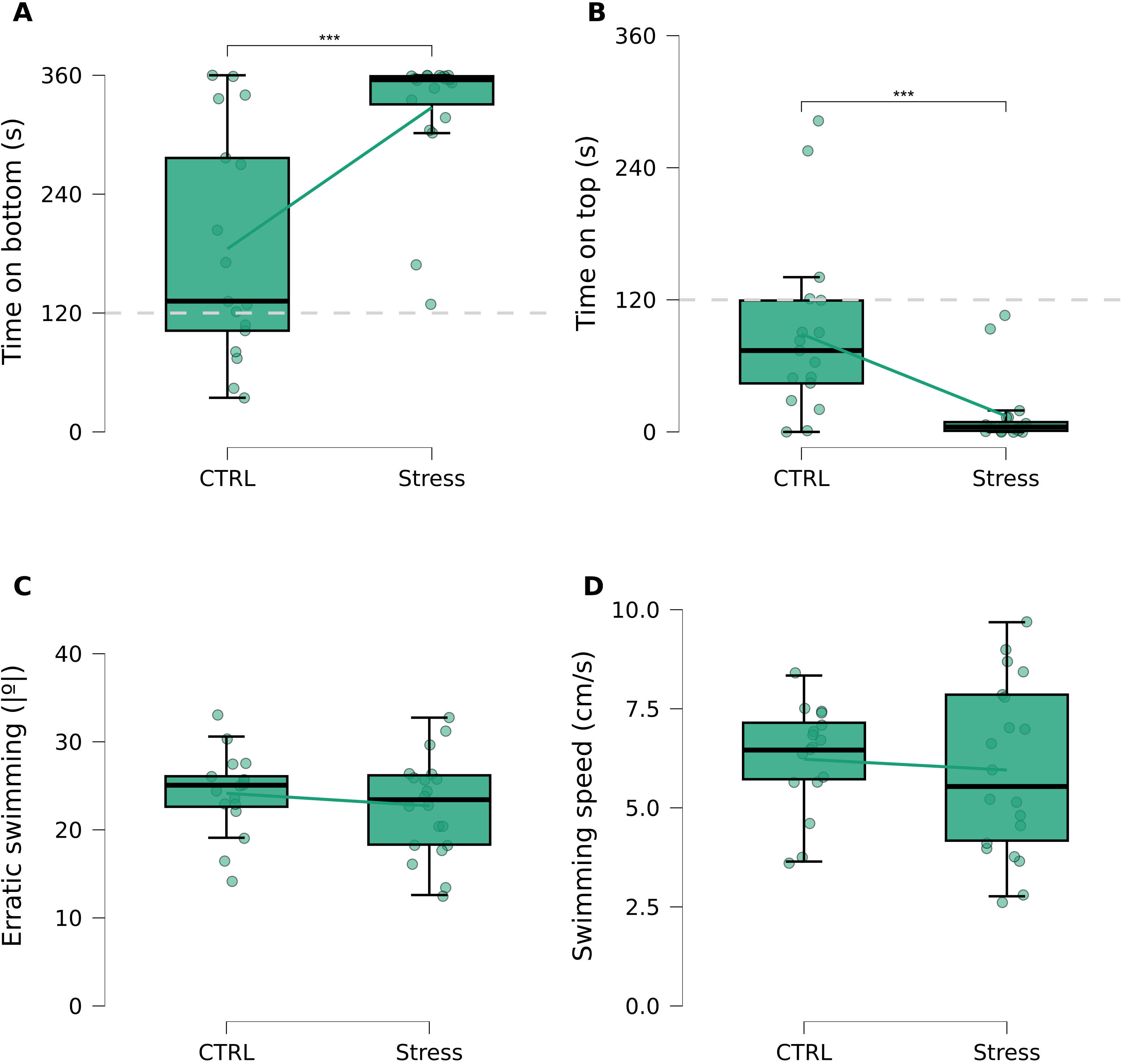
Unpredictable chronic stress (UCS) increases anxiety-like behavior in the zebrafish novel tank test. (A) Time on bottom. (B) Time on top. (C) Erratic swimming. (D) Swimming speed. Each dot represents a single individual, and boxplots represent median ± interquartile range. Lines connecting boxplots represent differences between group means. In (A) and (B), the dotted line (---) represents chance levels. For all panels, asterisks indicate significance levels: ***: p < 0.001.

A main effect of stage (SI vs. SN) was found for time near S1 (F_[1, 35]_ = 67.93, p < 0.001, η²p = 0.66); no main effects of stress were found for this variable (F_[1, 35]_= 0.1482, p = 0.703, η²p = 0.0042), but a stage X treatment interaction was also found (F_[1, 35]_ = 22.78, p < 0.001, η²p = 0.394). *Post-hoc* tests found that UCS decreased time near the conspecific during SI (t_[df= 35]_ = 2.981, p = 0.01, *d* = 1.095) and increased time near the “old stranger” during SN (t_[df= 35]_ = -4.393, p < 0.001, *d* = -1.263)(Figure 2A).

**Figure 2.**
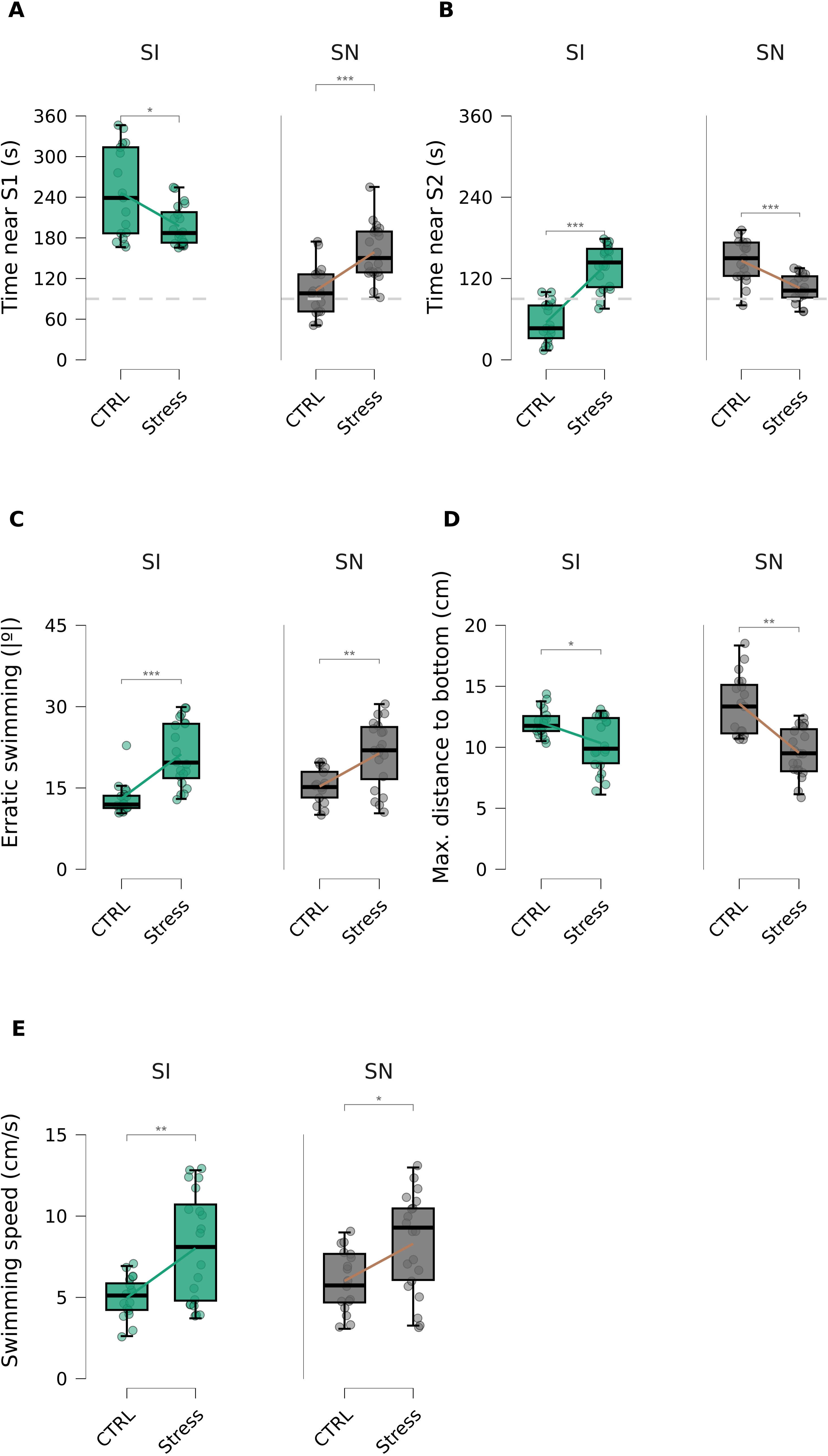
Unpredictable chronic stress (UCS) decreases social preference/increases social avoidance in the social preference test. (A) Time near conspecific (during SI) or “old stranger” (during SN). (B) Timer near empty tank (during SI) or “new stranger” (during SI). (C) Erratic swimming. (D) Maximum distance to tank bottom. (E) Swimming speed. For all panels, SI represents behavior during the social investigation stage, and SN represents behavior during the social novelty stage. Each dot represents a single individual, and boxplots represent median ± interquartile range. Lines connecting boxplots represent differences between group means. In (A) and (B), the dotted line (---) represents chance levels. For all panels, asterisks indicate significance levels: ***: p < 0.001; **: p < 0.01; *: p < 0.05.

A main effect of stage (SI vs. SN) was found for time near S2 (F_[1, 35]_ = 19.8, p < 0.001, η²p = 0.3614), and a main effect of stress was found (F_[1, 35]_ = 10.17, p = 0.003, η²p = 0.225). A stage X treatment interaction was also found (F_[1, 35]_ = 87.78, p < 0.001, η²p = 0.7149). *Post-hoc* tests found that UCS increased time near the empty tank during SI (t_[1, 35]_ = -8.129, p < 0.001, η²p = -2.198) and decreased time near the “new stranger” during SN (t_[df= 35]_ = 4.758, p < 0.001, η²p = 1.418)(Figure 2B)

No significant main effects of stage (SI vs. SN) was found for erratic swimming (F_[1, 35]_ = 0.866, p = 0.358, η²p = 0.0241), but a main effect of stress was found (F_[1, 35]_ = 50.84, p < 0.001, η²p = 0.592); no interaction effects were observed (F_[1, 35]_ = 0.6651, p = 0.425, η²p = 0.018). *Post-hoc* tests found that UCS increased erratic swimming throughout both stages (t_[df= 35]_ = -7.131, p < 0.001, *d* = -1.46)(Figure 2C).

No significant main effects of stage (SI vs. SN) were found for geotaxis (F_[1, 35]_ = 0.8513, p = 0.363, η²p = 0.024), but a main effect of stress was found (F_[1, 35]_ = 30.05, p < 0.001, η²p = 0.462). A stage X stress effect was also found (F_[1, 35]_ = 8.183, p = 0.007, η²p = 0.189). *Post-hoc* tests found that UCS decreased distance to bottom in both SI (t_[df= 35]_ = 2.289, p = 0.023, *d* = 0.8381) and SN (t_[df= 35]_ = 5.563, p < 0.001, η²p = 2.005)(Figure 2D); moreover, in control animals distance to bottom increased in SN in relation to SI (t_[df= 35]_ = -2.573, p = 0.029, *d* = -0.776).

No significant main effects of stage (SI vs. SN) were found for swimming speed (F_[1, 35]_ = 1.291, p = 0.264, η²p = 0.0336), but a main effect of stress was found (F_[1, 35]_ = 17.71, p < 0.001, η²p = 0.336); no significant interactions were found (F_[1, 35]_ = 0.431, p = 0.516, η²p = 0.012). *Post-hoc* tests found that UCS increased swimming speed throughout both stages (t_[df= 35]_ = -4.208, p < 0.001, *d* = -1.023)(Figure 2E).

For the TBARS assay, one brain sample and two heady kidney samples from the control group, and two brain samples and two head kidney samples from the UCS group were removed from the analysis because values were below the limit of quantification. An independent-samples Student’s *t*-test revealed a statistically significant difference between the control and UCS groups in brain tissue (t_[df= 32]_ = −8.154, p < 0.001, *d* = -2.802), indicating that the UCS group exhibited higher TBARS levels than the control group (Figure 3A). Similarly, UCS increased TBARS levels in the head kidney (t_[df= 31]_ = -3.357, p = 0.002, *d* = -1.174)(Figure 3B). When correlations between behavioral variables in the NTT and lipid peroxidation were analyzed, no significant correlation was found between brain or head kidney TBARS levels and any behavioral variables for the control group, but a significant correlation was found between brain TBARS and both time on bottom and time on top for the UCS group (Figures 3C and 3D). No significant correlations were found between TBARS levels and any behavioral variables for the control or UCS groups in SPT (Figures 3E and 3F).

**Figure 3.**
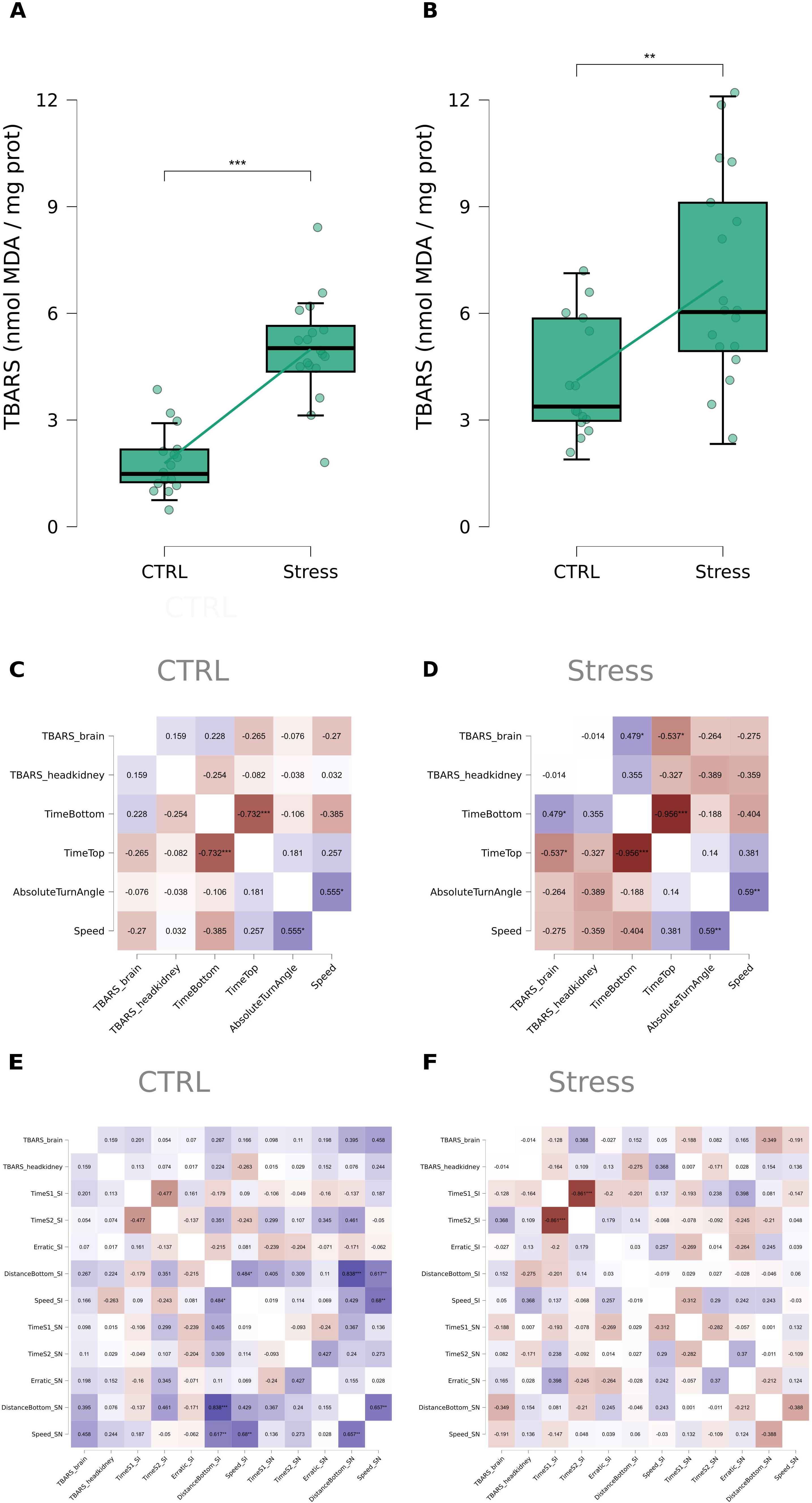
Unpredictable chronic stress (UCS) increases lipid peroxidation in brain and head kidney samples. (A) TBARS levels in brain tissues. (B) TBARS levels in head kidney tissues. (C) Correlations between sample TBARS levels and behavioral endpoints in the novel tank test (NTT) in control (CTRL) animals. (D) Correlations between sample TBARS levels and behavioral endpoints in the novel tank test in stressed (UCS) animals. (E) Correlations between sample TBARS levels and behavioral endpoints in the social preference test in control (CTRL) animals. (F) Correlations between sample TBARS levels and behavioral endpoints in the social preference test in stressed (UCS) animals. In (A) and (B), each dot represents a single individual, and boxplots represent median ± interquartile range. Lines connecting boxplots represent differences between group means. In (C-F), colors represent the magnitude and direction of the correlation, with red representing negative correlations and blue representing positive correlations. For all panels, asterisks indicate significance levels: ***: p < 0.001; **: p < 0.01; *: p < 0.05.

## 4. Discussion

The present work described the effects of unpredictable chronic stress (UCS), a paradigm that is used in rodents to model depressive-like phenotypes (Willner, 2017), in anxiety-like behavior in the zebrafish novel tank test (NTT), as well as in social motivation / withdrawal in a two-stage zebrafish social preference test (SPT), and associate these effects with changes in lipid peroxidation in brain and head kidney tissues. It was found that UCS increases anxiety-like behavior in the NTT, replicating previous work that showed similar effects (Gallas-Lopes et al., 2023). Moreover, it was found that UCS decreases social preference for a conspecific in the social investigation stage, increased preference for the known conspecific in the social novelty stage, and decreased preference for an unknown conspecific in that latter stage. While UCS also increased lipid peroxidation in the brains and head kidneys of stressed individuals, a correlation between peroxidation levels in the brains (but not head kidneys) and behavior was observed only in the NTT in stressed individuals.

The UCS paradigm has demonstrated reliability in inducing stress-related behavioral phenotype in zebrafish in the dimension of anxiety-like behavior, but so far effects on social behavior have been elusive (Gallas-Lopes et al., 2023; Neves et al., 2026). Our present results are consistent with findings demonstrating anxiogenic-like effects of UCS across behavioral tests in zebrafish (Gallas-Lopes et al., 2023), as well as on oxidative stress parameters (Lima-Maximino et al., 2025; Marcon et al., 2018; Mocelin et al., 2019); they also contribute to indirectly link brain oxidative stress to the extreme geotaxis observed in animals subjected to UCS by showing a positive correlation between these variables in stressed individuals.

The main novelty of the present study, however, lies in the demonstration that UCS disrupts social preference in zebrafish. A previous meta-analysis (Gallas-Lopes et al., 2023) failed to find significant effects of UCS on zebrafish social behavior across studies; however, not only heterogeneity was considerable, but the number of studies that used social preference were relatively small. Using a paradigm in which social approach responses were induced by showing videos of shoaling animals, Fulcher et al. (2017) did not find an effect of UCS on distance to the stimulus; nonetheless, unstressed animals also did not show to respond strongly to this stimulus, which makes it difficult to interpret the results. Reddy et al. (2021; 2018, 2019) exposed animals to a shorter period of UCS (7 days), and used a one-stage protocol for social preference, reporting that UCS decreased time in the interaction zone. Bertelli et al. (2021) and Neves et al. (2026) used a 14-day UCS protocol and did not observe changes in social preference in a one-stage SPT. Thus, significant differences in stress protocol (e.g., duration) and, crucially, in the SPT protocol exist between the present study and other studies in the literature.

The two-stage version of SPT offers a distinct advantage for evaluating depressive-like phenotypes because it dissociates social motivation from social novelty seeking. Stage 1 (social investigation) primarily captures social withdrawal, a core depression-related alteration, by measuring preference for a conspecific versus an empty compartment. Stage 2 (social novelty) then determines whether reduced sociability reflects a global loss of social interest or a more specific change in social valuation. This is important because if stressed fish still prefer the familiar conspecific, non-specific confounds such as locomotor impairment, sensory deficits, or generalized social aversion can be largely discarded. Thus, the two-stage design provides a more specific depression-related endpoint while also allowing the detection of anxiety-, neophobia-, or anhedonia-driven shifts in social novelty preference.

The present findings contrast with those of Bertelli et al. (2021) and Neves et al. (2026) primarily from a methodological standpoint. The present two-stage protocol detected reduced social motivation in Stage 1 (SI) and a selective increase in preference for the familiar conspecific in Stage 2 (SN), indicating that social interest was not globally abolished but rather shifted away from novel social cues. The most salient methodological difference is the social stimulus: Bertelli et al. (2021) and Neves et al. (2026) used a shoal of 10 conspecifics, whereas we used a single conspecific. A 10-fish stimulus likely produces a stronger, more unambiguous social signal that may be resistant to stress-induced reductions in social motivation (do Nascimento & Maximino, 2023), thereby masking an effect. In contrast, a single unfamiliar conspecific may be weaker, making the test more sensitive to stress-related social withdrawal or neophobia. Differences in stressor composition may also contribute, although both protocols produced anxiety-like effects in the novel tank test, indicating that both are effective stressors. Given the high statistical power in Neves et al. (2026), their result likely reflects a true difference in stimulus sensitivity rather than a Type II error. Therefore, our decreased preference in SI should not be interpreted as contradicting their findings; instead, it suggests that chronic stress reduces approach to a single conspecific, whereas attraction to a larger shoal may remain intact. Further studies comparing the effects of shoal size and other indices of stimulus salience are needed to untangle this hypothesis.

We interpret findings from our results in the SPT as consistent with a depressive-like phenotype. Differently from lab rodents, which usually show social avoidance of novel conspecifics (File, 2001; File & Hyde, 1978), zebrafish show a preference for conspecifics (social approach), preferring to investigate unfamiliar individuals (Barba-Escobedo & Gould, 2012; do Nascimento & Maximino, 2023; Engeszer et al., 2004; Ogi et al., 2021). Other studies suggest that visual contact with conspecifics is rewarding for zebrafish (Al-Imari & Gerlai, 2008; Scerbina et al., 2012), suggesting that zebrafish are motivated to investigate other individuals. The reduced preference for the conspecific in the social investigation phase indicates social withdrawal and diminished social motivation, which we suggest represent a core depression-like feature in zebrafish. During social novelty testing, the increased preference for the known conspecific and decreased preference for the unknown conspecific suggest that social recognition remains intact but that novelty seeking is impaired. This shift toward the familiar conspecific likely reflects heightened anxiety/neophobia or social anhedonia rather than a social memory deficit (de Abreu et al., 2022; Lai et al., 2023). Thus, unpredictable chronic stress appears to preserve social discrimination while reducing the incentive value of novel social interaction and increasing avoidance of unfamiliar social stimuli.

In addition to showing signs of social withdrawal / anhedonia in the SPT, other behavioral effects were also observed in this test. Specifically, stressed animals showed increased erratic swimming and geotaxis irrespective of the stage; this is consistent with findings from the NTT. Incidentally, no correlations were found between these variables and time near any of the conspecifics in any stage, suggesting that social motivation and anxiety-like behavior are dissociable in this test. Further studies, with larger samples that allow for the unbiased use of multivariate statistics, can better characterize this.

The TBARS analysis showed that UCS was associated with increased oxidative damage resulting from lipid peroxidation. In this regard, the increase in reactive oxygen species (ROS) suggests that exposure to stressors affects oxidative homeostasis, as indicated by the biochemical parameters evaluated in the brains and head kidneys of zebrafish. These results replicate previous findings that show that UCS increases oxidative stress in the zebrafish brain (Lima-Maximino et al., 2025; Marcon et al., 2018, 2019; Mocelin et al., 2019) and extend it to show oxidative effects in the head kidney, in which steroidogenic cells that produce cortisol and catecholamines are located (Alsop & Vijayan, 2009). The meta-analysis by Gallas-Lopes et al. (2023) showed that UCS consistently increases cortisol in zebrafish, and the effects on oxidative stress in the head kidney might reflect this. Nonetheless, TBARS levels in the head kidney did not correlate with changes in behavior, contrary to TBARS levels in the brain. This latter result also suggests a causal link between brain oxidative stress and the anxiogenic-like effects of UCS in zebrafish, an hypothesis further supported by findings that antioxidants rescue the effects of UCS (Mocelin et al., 2019).

The findings of Marcon et al. (2019) and Mocelin et al. (2019) further support the understanding that chronic stress has a significant impact on redox homeostasis by compromising endogenous antioxidant defense systems. They observed an increase in lipid peroxidation that was accompanied by decreased non-protein thiol and impaired superoxide dismutase activities, suggesting that UCS intensifies the accumulation of reactive oxygen species (ROS), especially superoxide, through a defect in superoxide scavenging systems. The findings of the present study suggest that an imbalance between the production and removal of free radicals may represent a mechanism mediating neuronal damage triggered by prolonged stress. Other findings from the literature suggest that superoxide accumulation and glutathione depletion resulting from UCS lead to upregulation of mitochondrial proteins that try to compensate for the results of oxidative stress, ultimately failing (Chakravarty et al., 2013), fueling an inflammatory NF-κB/n-6 PUFA feedforward loop (Kotova et al., 2025; Pinto et al., 2026; B. R. Reddy et al., 2021) that produces even more oxidative stress. One important consequence is a change in cell proliferation in the telencephalon, increasing it in the dorsomedial zone (homologous to the associative amygdala) and decreasing it in the dorsolateral zone (homologous to the hippocampus)(B. R. Reddy et al., 2021). Thus, these data contribute to the understanding of how oxidative imbalance may act as one of the major pathophysiological factors associated with the behavioral alterations observed in animal models exposed to persistent stressors (Che et al., 2015).

Although the behavioral outcomes reported across studies may differ partially, the available evidence indicates that unpredictable chronic stress produces behavioral effects in zebrafish that are consistent with a mixed anxiogenic/depressive-like profile. Therefore, UCS may represent a valuable model for investigating stress-related mechanisms underlying psychological disorders such as anxiety and depression. In this context, further methodological standardization is needed to establish the most appropriate experimental approach, given the variability in findings across studies, as well as differences in the methodologies used to implement UCS protocols.

## 5. Conclusion

The present study investigated the effects of an adapted unpredictable chronic stress (UCS) protocol (Piato et al., 2011) on depression-like behavioral parameters, particularly anhedonia, in zebrafish, an organism of considerable importance in behavioral, biochemical, and pharmacological research. Based on the results obtained, significant differences between the UCS and control groups were identified through the behavioral tests. In the novel tank test, significant differences were observed in the time spent at the top and bottom of the apparatus. In the social preference test, the animals showed differences in the time spent near the conspecific during the social interaction stage, as well as a social neophobic reaction in the social novelty stage. The biochemical findings also revealed significant alterations in TBARS levels, indicating increased lipid peroxidation and an association between exposure to stressors and oxidative imbalance. Based on these findings, it can be inferred that excessive exposure to stressful situations induces behavioral and biochemical changes in zebrafish that are consistent with an anxiogenic-like and depressogenic-like effect. Considering the results obtained in the present study and the existing evidence in the literature, further studies are needed to elucidate the relationship between the behavioral effects of chronic stress and the underlying neurochemical mechanisms involved in these alterations.

## Supplementary Files

**Supplementary File S1 -** Sex and morphometric baseline data for the animals used in the experiments.

**Supplementary File S2 -** Data and statistical analyses considering sex as an independent variable.

## Notes

### Competing Interest Statement

The authors have declared no competing interest.

### Funding

- Fundação Amazônia de Amparo a Estudos e Pesquisas do Pará: Termo de Outorga nº 050/2021

