## Supplementary File S2 for "Unpredictable chronic stress produces a mixed depression/anxiety behavioral profile in zebrafish"

#### Gráficos

##### TimeBottom

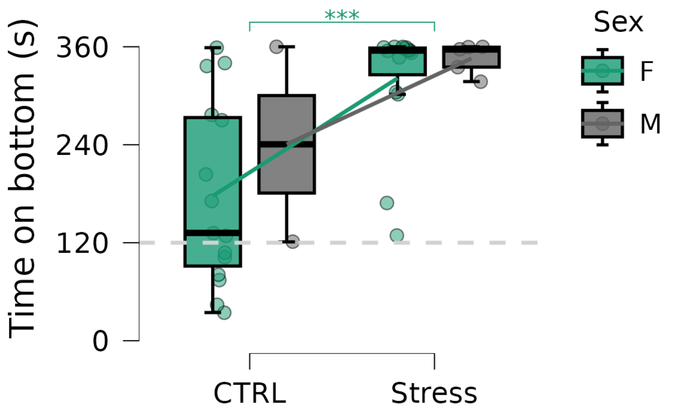


##### TimeTop

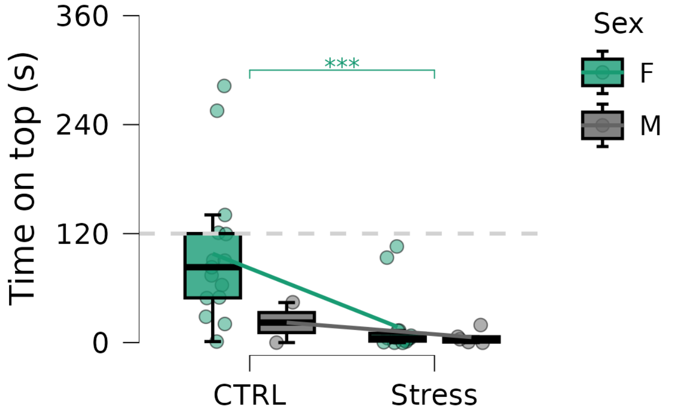

**Erratic**

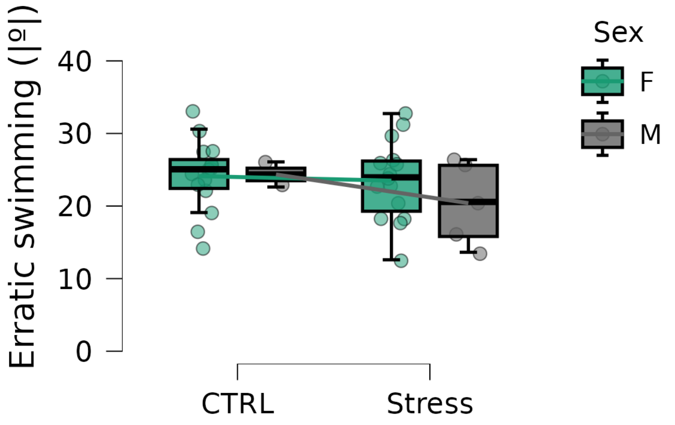


**Speed**

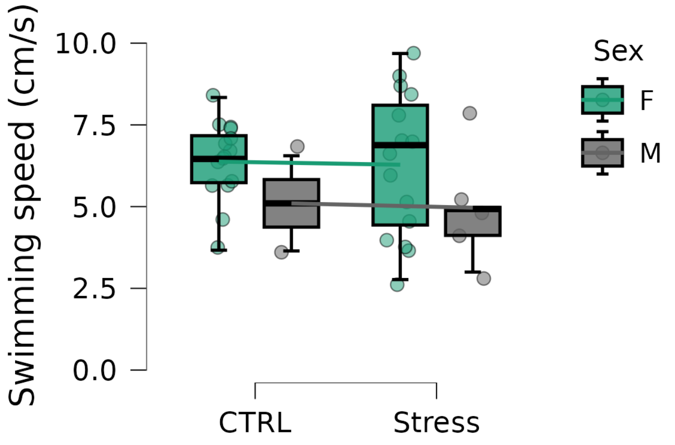


#### IDs de gráficos disponíveis

Available Plot IDs: TimeBottom, TimeTop, Erratic, Speed

**Gráfico Grid**

**Combined Plot Grid**

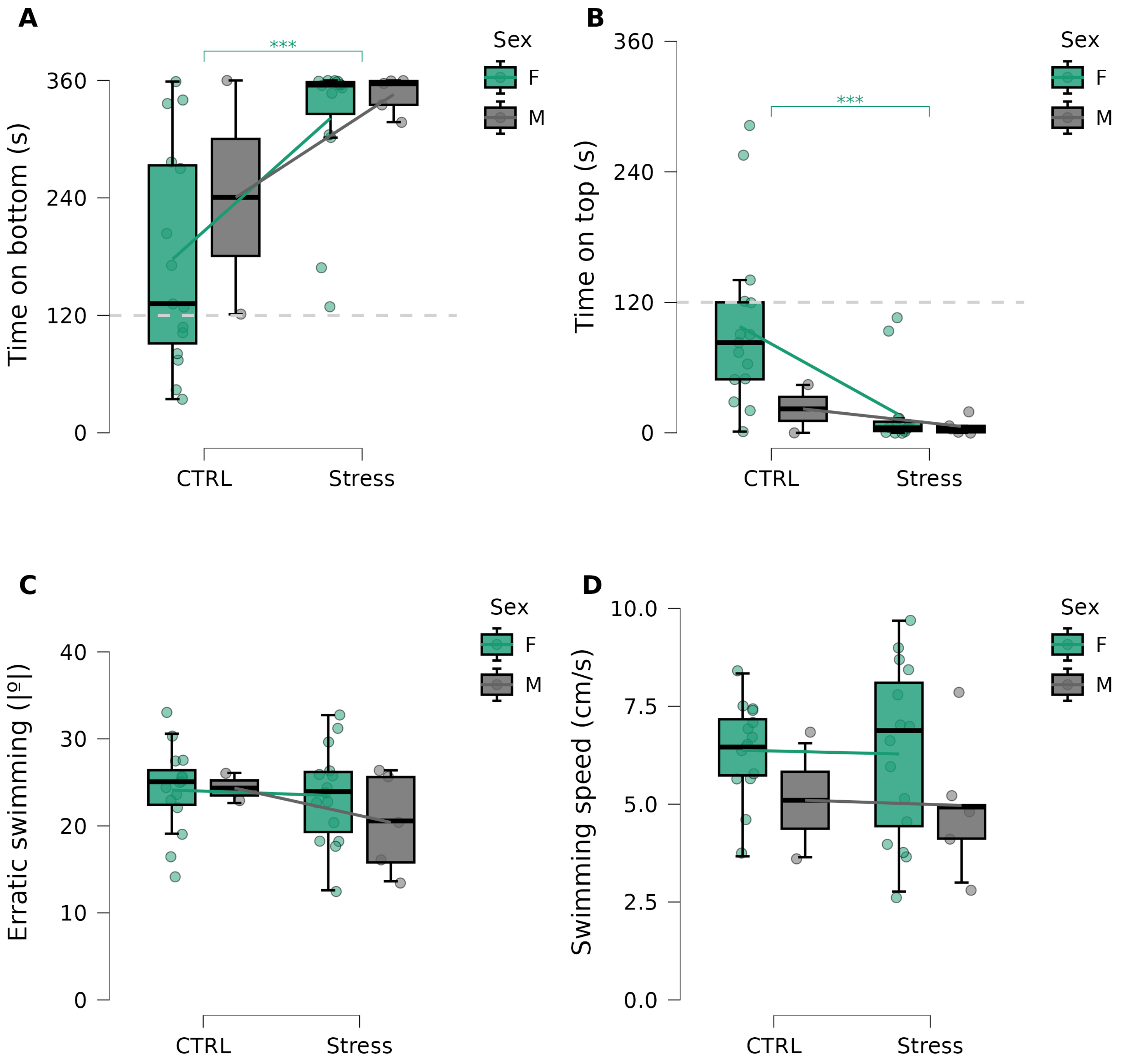

### TBARS

#### Gráficos

##### Brain

##### Head kidney

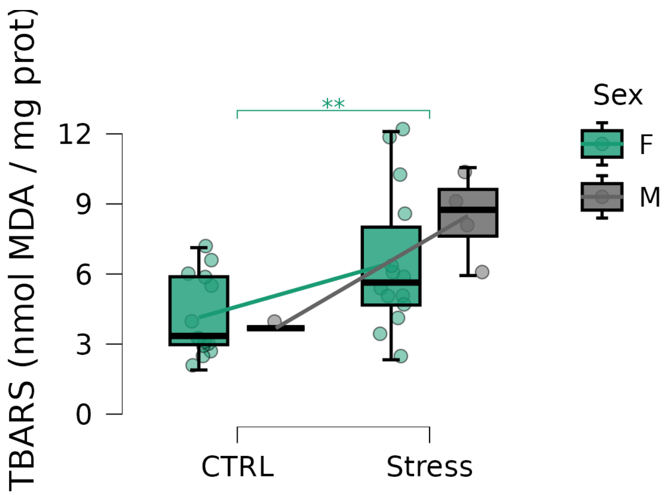


#### IDs de gráficos disponíveis

Available Plot IDs: Brain, Head kidney

Combined Plot Grid

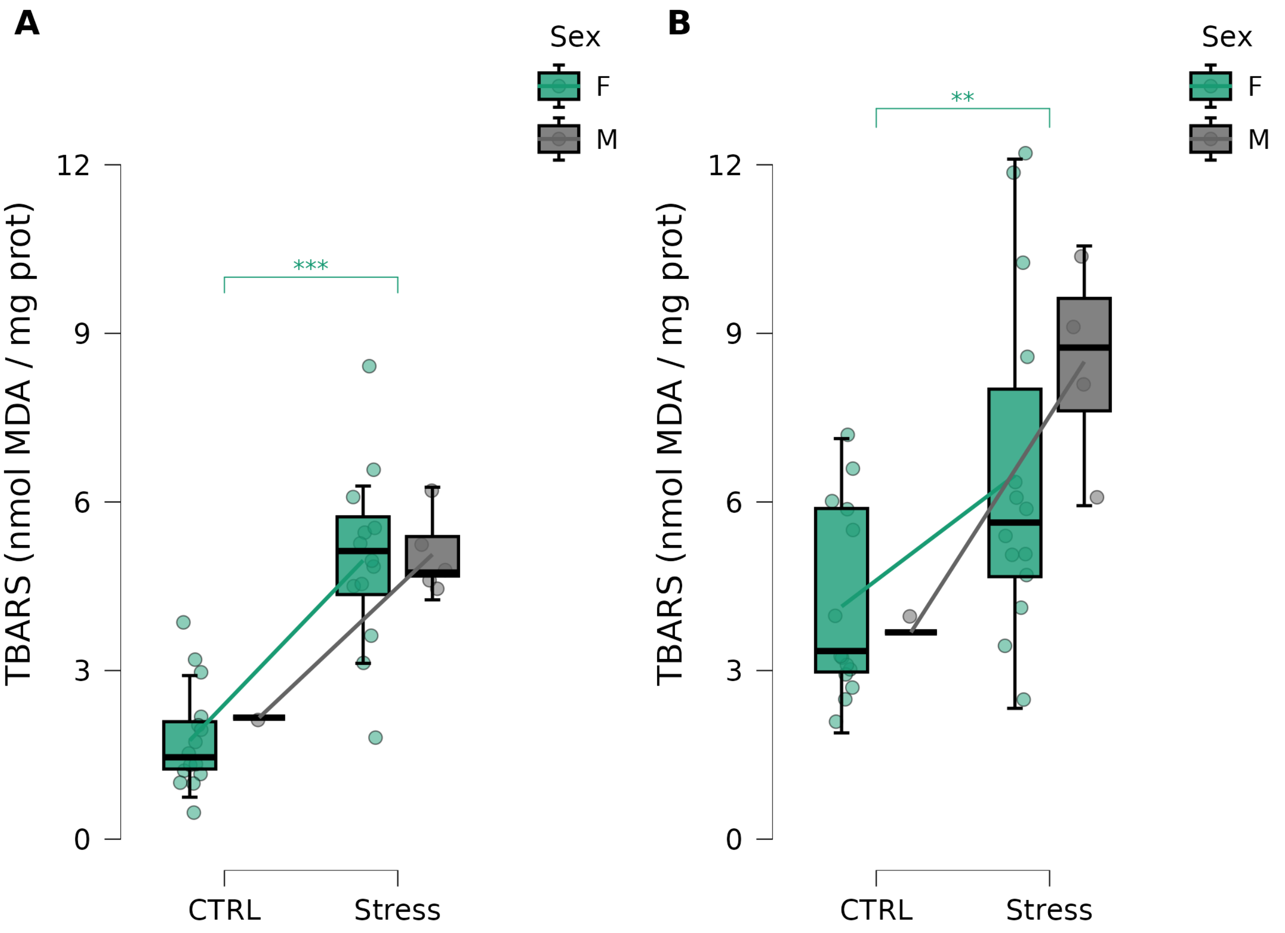

### ANOVA - Time on bottom

*ANOVA - TimeBottom*

| Casos | Soma dos quadrados | gl | Média Quadrática | F | p | $\eta^2_p$ |
|---|---|---|---|---|---|---|
| Sex | 9178 | 1 | 9178 | 1.076 | .307 | 0.03157 |
| Group | $7.459\times10^{+4}$ | 1 | $7.459\times10^{+4}$ | 8.743 | .006 | 0.2094 |
| Sex ∗ Group | 1821 | 1 | 1821 | 0.2135 | .647 | 0.006428 |
| Residuals | $2.815\times10^{+5}$ | 33 | 8531 | | | |

*Nota.* Soma dos quadrados Tipo III

### Testes Post Hoc

#### Padrão (DHS)

*Comparações Post Hoc - Group*

| | | Diferença Média | EP | gl | t | D de Cohen | $p_{Tukey}$ |
|---|---|---|---|---|---|---|---|
| CTRL | Stress | −124.7 | 42.16 | 33 | −2.957 | −1.350 | .006** |

** p < .01

*Nota.* Os resultados são calculados em média nos níveis de:Sex

*Letter-Based Grouping - Group*

| Group | Letter |
|---|---|
| CTRL | a |
| Stress | b |

*Nota.* If two or more means share the same grouping symbol,
then we cannot show them to be different, but we also did not show them to be the same.

*Comparações Post Hoc - Sex ✻ Group*

| | | Diferença Média | EP | gl | t | D de Cohen | $p_{Tukey}$ |
|---|---|---|---|---|---|---|---|
| F CTRL | M CTRL | −63.21 | 69.53 | 33 | −0.9091 | −0.6843 | .800 |
| | F Stress | −144.1 | 33.73 | 33 | −4.274 | −1.561 | < .001*** |
| | M Stress | −168.4 | 47.70 | 33 | −3.530 | −1.823 | .007** |
| M CTRL | F Stress | −80.93 | 69.53 | 33 | −1.164 | −0.8762 | .653 |
| | M Stress | −105.2 | 77.28 | 33 | −1.361 | −1.139 | .532 |
| F Stress | M Stress | −24.25 | 47.70 | 33 | −0.5084 | −0.2625 | .957 |

** p < .01, *** p < .001

*Nota.* Valor-p ajustado para comparar uma família de 4 estimativas.

*Letter-Based Grouping - Sex ✻ Group*

| Sex | Group | Letter |
|---|---|---|
| F | CTRL | a |
| M | | ab |
| F | Stress | b |
| M | | b |

*Nota.* If two or more means share the same grouping symbol,
then we cannot show them to be different, but we also did not show them to be the same.

### ANOVA - Time on top

*ANOVA - TimeTop*

| Casos | Soma dos quadrados | gl | Média Quadrática | F | p | $\eta^2_p$ |
|---|---|---|---|---|---|---|
| Sex | 9127 | 1 | 9127 | 2.843 | .101 | 0.07933 |
| Group | $1.120\times10^{+4}$ | 1 | $1.120\times10^{+4}$ | 3.489 | .071 | 0.09562 |
| Sex ∗ Group | 4985 | 1 | 4985 | 1.553 | .221 | 0.04495 |
| Residuals | $1.059\times10^{+5}$ | 33 | 3210 | | | |

*Nota.* Soma dos quadrados Tipo III

### Testes Post Hoc

#### Padrão (DHS)

*Comparações Post Hoc - Group*

| | | Diferença Média | EP | gl | t | D de Cohen | $p_{Tukey}$ |
|---|---|---|---|---|---|---|---|
| CTRL | Stress | 48.30 | 25.86 | 33 | 1.868 | 0.8526 | .071 |

*Nota.* Os resultados são calculados em média nos níveis de:Sex

*Letter-Based Grouping - Group*

| Group | Letter |
|---|---|
| CTRL | a |
| Stress | a |

*Nota.* If two or more means share the same grouping symbol,
then we cannot show them to be different, but we also did not show them to be the same.

*Comparações Post Hoc - Sex ✻ Group*

| | | Diferença Média | EP | gl | t | D de Cohen | $p_{Tukey}$ |
|---|---|---|---|---|---|---|---|
| F CTRL | M CTRL | 75.83 | 42.65 | 33 | 1.778 | 1.338 | .302 |
| | F Stress | 80.53 | 20.69 | 33 | 3.893 | 1.421 | .002** |
| | M Stress | 91.91 | 29.26 | 33 | 3.141 | 1.622 | .018* |
| M CTRL | F Stress | 4.697 | 42.65 | 33 | 0.1101 | 0.08291 | 1.000 |
| | M Stress | 16.08 | 47.40 | 33 | 0.3391 | 0.2838 | .986 |
| F Stress | M Stress | 11.38 | 29.26 | 33 | 0.3889 | 0.2008 | .980 |

* $p < .05$, ** $p < .01$

*Nota.* Valor-p ajustado para comparar uma família de 4 estimativas.

*Letter-Based Grouping - Sex ✻ Group*

| Sex | Group | Letter |
|---|---|---|
| F | CTRL | b |
| M | | ab |
| F | Stress | a |
| M | | a |

*Nota.* If two or more means share the same grouping symbol,
then we cannot show them to be different, but we also did not show them to be the same.

### ANOVA - Erratic swimming

*ANOVA - AbsoluteTurnAngle*

| Casos | Soma dos quadrados | gl | Média Quadrática | F | p | $\eta^2_p$ |
|---|---|---|---|---|---|---|
| Sex | 9.880 | 1 | 9.880 | 0.3642 | .550 | 0.01092 |
| Group | 25.11 | 1 | 25.11 | 0.9258 | .343 | 0.02729 |
| Sex ∗ Group | 13.32 | 1 | 13.32 | 0.4909 | .488 | 0.01466 |
| Residuals | 895.1 | 33 | 27.13 | | | |

*Nota.* Soma dos quadrados Tipo III

### Testes Post Hoc

#### Padrão (DHS)

*Comparações Post Hoc - Group*

| | | Diferença Média | EP | gl | t | D de Cohen | $p_{Tukey}$ |
|---|---|---|---|---|---|---|---|
| CTRL | Stress | 2.287 | 2.377 | 33 | 0.9622 | 0.4392 | .343 |

*Nota.* Os resultados são calculados em média nos níveis de:Sex

*Letter-Based Grouping - Group*

| Group | Letter |
|---|---|
| CTRL | a |
| Stress | a |

*Nota.* If two or more means share the same grouping symbol,
then we cannot show them to be different, but we also did not show them to be the same.

*Comparações Post Hoc - Sex ✻ Group*

| | | Diferença Média | EP | gl | t | D de Cohen | $p_{Tukey}$ |
|---|---|---|---|---|---|---|---|
| F CTRL | M CTRL | −0.2309 | 3.921 | 33 | −0.05890 | −0.04434 | 1.000 |
| | F Stress | 0.6217 | 1.902 | 33 | 0.3269 | 0.1194 | .988 |
| | M Stress | 3.722 | 2.689 | 33 | 1.384 | 0.7146 | .518 |
| M CTRL | F Stress | 0.8526 | 3.921 | 33 | 0.2175 | 0.1637 | .996 |
| | M Stress | 3.953 | 4.357 | 33 | 0.9072 | 0.7590 | .801 |
| F Stress | M Stress | 3.100 | 2.689 | 33 | 1.153 | 0.5953 | .660 |

*Nota.* Valor-p ajustado para comparar uma família de 4 estimativas.

*Letter-Based Grouping - Sex ✻ Group*

| Sex | Group | Letter |
|---|---|---|
| F | CTRL | a |
| M | | a |
| F | Stress | a |
| M | | a |

*Nota.* If two or more means share the same grouping symbol,
then we cannot show them to be different, but we also did not show them to be the same.

### ANOVA - Swimming speed

*ANOVA - Speed*

| Casos | Soma dos quadrados | gl | Média Quadrática | F | p | $\eta^2_p$ |
|---|---|---|---|---|---|---|
| Sex | 8.056 | 1 | 8.056 | 2.613 | .116 | 0.07337 |
| Group | 0.06221 | 1 | 0.06221 | 0.02018 | .888 | $6.110 \times 10^{-4}$ |
| Sex ∗ Group | 0.001725 | 1 | 0.001725 | $5.594 \times 10^{-4}$ | .981 | $1.695 \times 10^{-5}$ |
| Residuals | 101.7 | 33 | 3.083 | | | |

*Nota.* Soma dos quadrados Tipo III

### Testes Post Hoc

#### Padrão (DHS)

*Comparações Post Hoc - Group*

| | | Diferença Média | EP | gl | t | D de Cohen | $p_{Tukey}$ |
|---|---|---|---|---|---|---|---|
| CTRL | Stress | 0.1138 | 0.8015 | 33 | 0.1420 | 0.06483 | .888 |

*Nota.* Os resultados são calculados em média nos níveis de:Sex

*Letter-Based Grouping - Group*

| Group | Letter |
|---|---|
| CTRL | a |
| Stress | a |

*Nota.* If two or more means share the same grouping symbol,
then we cannot show them to be different, but we also did not show them to be the same.

*Comparações Post Hoc - Sex ✻ Group*

| | | Diferença Média | EP | gl | t | D de Cohen | $p_{Tukey}$ |
|---|---|---|---|---|---|---|---|
| F CTRL | M CTRL | 1.277 | 1.322 | 33 | 0.9658 | 0.7270 | .770 |
| | F Stress | 0.09489 | 0.6412 | 33 | 0.1480 | 0.05404 | .999 |
| | M Stress | 1.409 | 0.9067 | 33 | 1.554 | 0.8026 | .418 |
| M CTRL | F Stress | −1.182 | 1.322 | 33 | −0.8940 | −0.6730 | .808 |
| | M Stress | 0.1328 | 1.469 | 33 | 0.09039 | 0.07563 | 1.000 |
| F Stress | M Stress | 1.314 | 0.9067 | 33 | 1.450 | 0.7486 | .479 |

*Nota.* Valor-p ajustado para comparar uma família de 4 estimativas.

*Letter-Based Grouping - Sex ✻ Group*

| Sex | Group | Letter |
|---|---|---|
| F | CTRL | a |
| M | | a |
| F | Stress | a |
| M | | a |

*Nota.* If two or more means share the same grouping symbol,
then we cannot show them to be different, but we also did not show them to be the same.

### ANOVA - Brain TBARS

*ANOVA - TBARS_brain*

| Casos | Soma dos quadrados | gl | Média Quadrática | F | p | $\eta^2_p$ |
|---|---|---|---|---|---|---|
| Sex | 0.1956 | 1 | 0.1956 | 0.1406 | .710 | 0.004664 |
| Group | 27.76 | 1 | 27.76 | 19.95 | < .001 | 0.3994 |
| Sex ∗ Group | 0.06610 | 1 | 0.06610 | 0.04751 | .829 | 0.001581 |
| Residuals | 41.74 | 30 | 1.391 | | | |

*Nota.* Soma dos quadrados Tipo III

#### Testes Post Hoc

##### Padrão (DHS)

*Comparações Post Hoc - Group*

| | | Diferença Média | EP | gl | t | D de Cohen | $p_{Tukey}$ |
|---|---|---|---|---|---|---|---|
| CTRL | Stress | −3.053 | 0.6836 | 30 | −4.467 | −2.589 | < .001*** |

*** p < .001

*Nota.* Os resultados são calculados em média nos níveis de:Sex

*Letter-Based Grouping - Group*

| Group | Letter |
|---|---|
| CTRL | a |
| Stress | b |

*Nota.* If two or more means share the same grouping symbol,
then we cannot show them to be different, but we also did not show them to be the same.

*Comparações Post Hoc - Sex ✻ Group*

| | | Diferença Média | EP | gl | t | D de Cohen | $p_{Tukey}$ |
|---|---|---|---|---|---|---|---|
| F CTRL | M CTRL | −0.4053 | 1.218 | 30 | −0.3327 | −0.3436 | .987 |
| | F Stress | −3.202 | 0.4469 | 30 | −7.165 | −2.715 | < .001*** |
| | M Stress | −3.310 | 0.6091 | 30 | −5.434 | −2.806 | < .001*** |
| M CTRL | F Stress | −2.797 | 1.224 | 30 | −2.285 | −2.371 | .124 |
| | M Stress | −2.904 | 1.292 | 30 | −2.248 | −2.462 | .134 |
| F Stress | M Stress | −0.1073 | 0.6207 | 30 | −0.1729 | −0.09098 | .998 |

*** p < .001

*Nota.* Valor-p ajustado para comparar uma família de 4 estimativas.

*Letter-Based Grouping - Sex ✻ Group*

| Sex | Group | Letter |
|---|---|---|
| F | CTRL | a |
| M | | ab |
| F | Stress | b |
| M | | b |

*Nota.* If two or more means share the same grouping symbol,
then we cannot show them to be different, but we also did not show them to be the same.

### ANOVA - Head kidney TBARS

*ANOVA - TBARS_headkidney*

| Casos | Soma dos quadrados | gl | Média Quadrática | F | p | $\eta^2_p$ |
|---|---|---|---|---|---|---|
| Sex | 1.754 | 1 | 1.754 | 0.3078 | .583 | 0.01050 |
| Group | 36.71 | 1 | 36.71 | 6.440 | .017 | 0.1817 |
| Sex ∗ Group | 4.425 | 1 | 4.425 | 0.7763 | .386 | 0.02607 |
| Residuals | 165.3 | 29 | 5.699 | | | |

*Nota.* Soma dos quadrados Tipo III

#### Testes Post Hoc

##### Padrão (DHS)

*Comparações Post Hoc - Group*

| | | Diferença Média | EP | gl | t | D de Cohen | $p_{Tukey}$ |
|---|---|---|---|---|---|---|---|
| CTRL | Stress | −3.575 | 1.409 | 29 | −2.538 | −1.498 | .017* |

* p < .05

*Nota.* Os resultados são calculados em média nos níveis de:Sex

*Letter-Based Grouping - Group*

| Group | Letter |
|---|---|
| CTRL | a |
| Stress | b |

*Nota.* If two or more means share the same grouping symbol,
then we cannot show them to be different, but we also did not show them to be the same.

*Comparações Post Hoc - Sex ✻ Group*

| | | Diferença Média | EP | gl | t | D de Cohen | $p_{Tukey}$ |
|---|---|---|---|---|---|---|---|
| F CTRL | M CTRL | 0.4597 | 2.471 | 29 | 0.1860 | 0.1925 | .998 |
| | F Stress | −2.334 | 0.9023 | 29 | −2.587 | −0.9776 | .068 |
| | M Stress | −4.357 | 1.354 | 29 | −3.219 | −1.825 | .016* |
| M CTRL | F Stress | −2.794 | 2.471 | 29 | −1.130 | −1.170 | .674 |
| | M Stress | −4.816 | 2.669 | 29 | −1.804 | −2.017 | .292 |
| F Stress | M Stress | −2.023 | 1.354 | 29 | −1.495 | −0.8473 | .454 |

* p < .05

*Nota.* Valor-p ajustado para comparar uma família de 4 estimativas.

*Letter-Based Grouping - Sex ✻ Group*

| Sex | Group | Letter |
|---|---|---|
| F | CTRL | a |
| M | | ab |
| F | Stress | ab |
| M | | b |

*Nota.* If two or more means share the same grouping symbol,
then we cannot show them to be different, but we also did not show them to be the same.

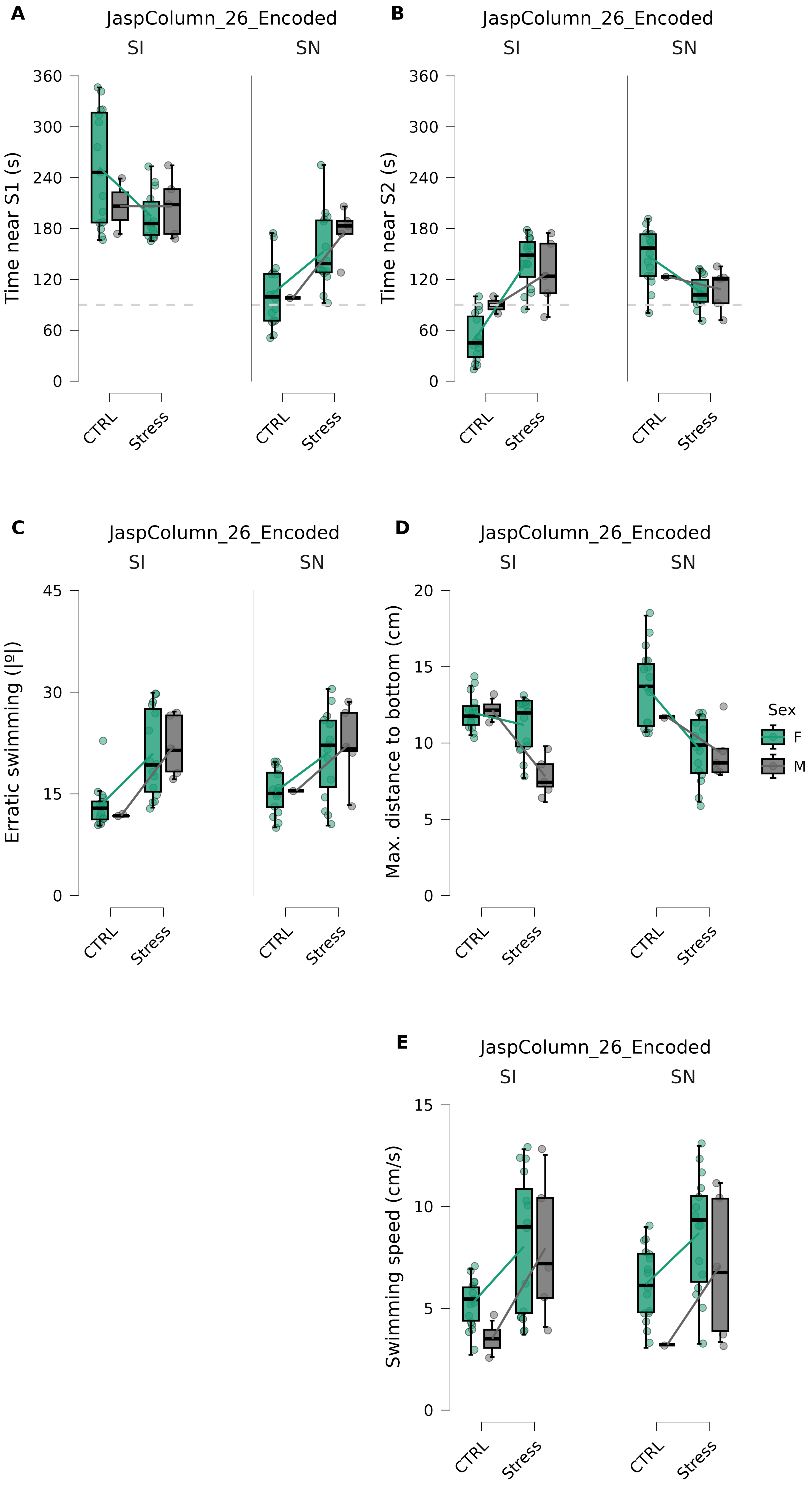

A
JaspColumn_26_Encoded
SI
SN
Time near S1 (s)
0
60
120
180
240
300
360
CTRL
Stress
B
JaspColumn_26_Encoded
SI
SN
Time near S2 (s)
0
60
120
180
240
300
360
CTRL
Stress
C
JaspColumn_26_Encoded
SI
SN
Erratic swimming (|º|)
0
15
30
45
CTRL
Stress
D
JaspColumn_26_Encoded
SI
SN
Max. distance to bottom (cm)
0
5
10
15
20
CTRL
Stress
Sex
F
M
E
JaspColumn_26_Encoded
SI
SN
Swimming speed (cm/s)
0
5
10
15
CTRL
Stress

### SPT

#### RMANOVA - Time near S1

*Within Subjects Effects*

| Cases | Sum of Squares | df | Mean Square | F | p | $\eta^2_p$ |
|---|---|---|---|---|---|---|
| SPT Stage | $5.313\times10^{+4}$ | 1 | $5.313\times10^{+4}$ | 24.06 | < .001 | 0.4217 |
| SPT Stage ∗ Treatment | $1.438\times10^{+4}$ | 1 | $1.438\times10^{+4}$ | 6.514 | .016 | 0.1648 |
| SPT Stage ∗ Sex | 5089 | 1 | 5089 | 2.305 | .138 | 0.06529 |
| SPT Stage ∗ Treatment ∗ Sex | 2963 | 1 | 2963 | 1.342 | .255 | 0.03908 |
| Residuals | $7.286\times10^{+4}$ | 33 | 2208 | | | |

*Note.* Type III Sum of Squares

*Between Subjects Effects*

| Cases | Sum of Squares | df | Mean Square | F | p | $\eta^2_p$ |
|---|---|---|---|---|---|---|
| Treatment | 949.6 | 1 | 949.6 | 0.5269 | .473 | 0.01572 |
| Sex | 366.1 | 1 | 366.1 | 0.2032 | .655 | 0.006119 |
| Treatment ∗ Sex | 1173 | 1 | 1173 | 0.6510 | .426 | 0.01934 |
| Residuals | $5.947\times10^{+4}$ | 33 | 1802 | | | |

*Note.* Type III Sum of Squares

#### Descriptives

##### Descriptives plots

###### SPT Stage: SI

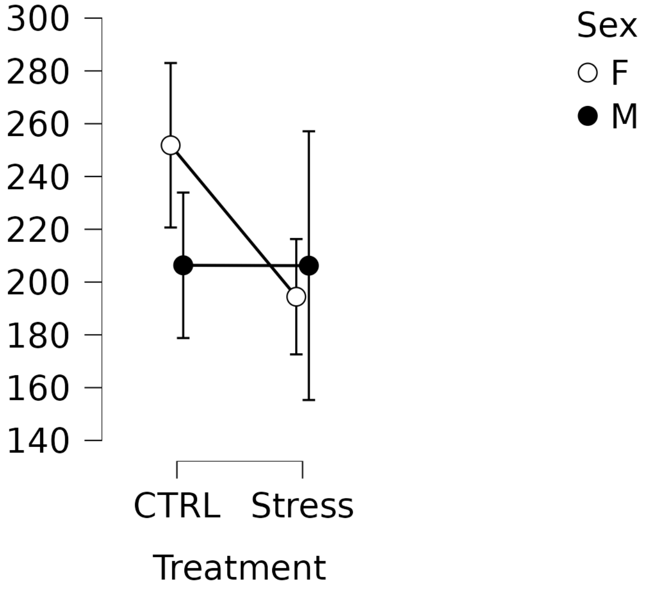


###### SPT Stage: SN

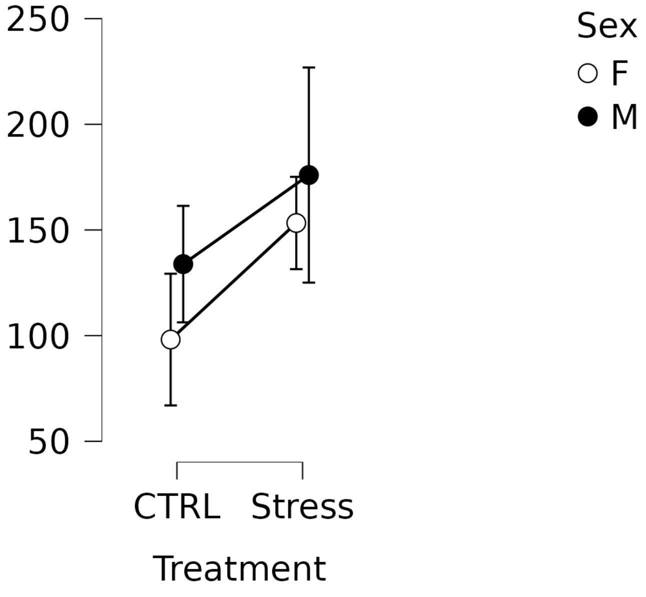

#### Post Hoc Tests

*Post Hoc Comparisons - Treatment*

| | | Mean Difference | SE | df | t | $p_{Holm}$ |
|---|---|---|---|---|---|---|
| CTRL | Stress | −9.946 | 13.70 | 33 | −0.7259 | .473 |

*Note.* Results are averaged over the levels of: Sex, SPT Stage

*Letter-Based Grouping - Treatment*

| Treatment | Letter |
|---|---|
| CTRL | a |
| Stress | a |

*Note.* If two or more means share the same grouping symbol, then we cannot show them to be different, but we also did not show them to be the same.

*Post Hoc Comparisons - SPT Stage*

| | | Mean Difference | SE | df | t | $p_{Holm}$ |
|---|---|---|---|---|---|---|
| SI | SN | 74.39 | 15.17 | 33 | 4.906 | < .001*** |

*** p < .001

*Note.* Results are averaged over the levels of: Treatment, Sex

*Letter-Based Grouping - SPT Stage*

| SPT Stage | Letter |
|---|---|
| SI | b |
| SN | a |

*Note.* If two or more means share the same grouping symbol, then we cannot show them to be different, but we also did not show them to be the same.

*Post Hoc Comparisons - Treatment ✻ SPT Stage*

| | | Mean Difference | SE | df | t | $p_{Holm}$ |
|---|---|---|---|---|---|---|
| CTRL, SI | Stress, SI | 28.76 | 22.90 | 33 | 1.256 | .218 |
| | CTRL, SN | 113.1 | 25.01 | 33 | 4.522 | < .001*** |
| | Stress, SN | 64.45 | 21.36 | 33 | 3.018 | .020* |
| Stress, SI | CTRL, SN | 84.34 | 19.48 | 33 | 4.330 | < .001*** |
| | Stress, SN | 35.69 | 17.16 | 33 | 2.080 | .091 |
| CTRL, SN | Stress, SN | −48.65 | 17.64 | 33 | −2.758 | .028* |

* p < .05, *** p < .001

*Note.* P-value adjusted for comparing a family of 6 estimates.

*Note.* Results are averaged over the levels of: Sex

*Letter-Based Grouping - Treatment:SPT Stage*

| Treatment | SPT Stage | Letter |
|---|---|---|
| CTRL | SI | c |
| Stress | | bc |
| CTRL | SN | a |
| Stress | | b |

*Note.* If two or more means share the same grouping symbol, then we cannot show them to be different, but we also did not show them to be the same.

### RMANOVA - Time near S2

*Within Subjects Effects*

| Cases | Sum of Squares | df | Mean Square | F | p | $\eta^2_p$ |
|---|---|---|---|---|---|---|
| SPT Stage | 5352 | 1 | 5352 | 6.813 | .014 | 0.1711 |
| SPT Stage ∗ Treatment | $2.573 \times 10^{+4}$ | 1 | $2.573 \times 10^{+4}$ | 32.75 | < .001 | 0.4981 |
| SPT Stage ∗ Sex | 347.4 | 1 | 347.4 | 0.4422 | .511 | 0.01322 |
| SPT Stage ∗ Treatment ∗ Sex | 2039 | 1 | 2039 | 2.595 | .117 | 0.07291 |
| Residuals | $2.592 \times 10^{+4}$ | 33 | 785.6 | | | |

*Note.* Type III Sum of Squares

*Between Subjects Effects*

| Cases | Sum of Squares | df | Mean Square | F | p | $\eta^2_p$ |
|---|---|---|---|---|---|---|
| Treatment | 1611 | 1 | 1611 | 1.936 | .173 | 0.05542 |
| Sex | 421.6 | 1 | 421.6 | 0.5066 | .482 | 0.01512 |
| Treatment ∗ Sex | 1364 | 1 | 1364 | 1.639 | .209 | 0.04733 |
| Residuals | $2.746 \times 10^{+4}$ | 33 | 832.2 | | | |

*Note.* Type III Sum of Squares

#### Descriptives

##### Descriptives plots

###### SPT Stage: SI

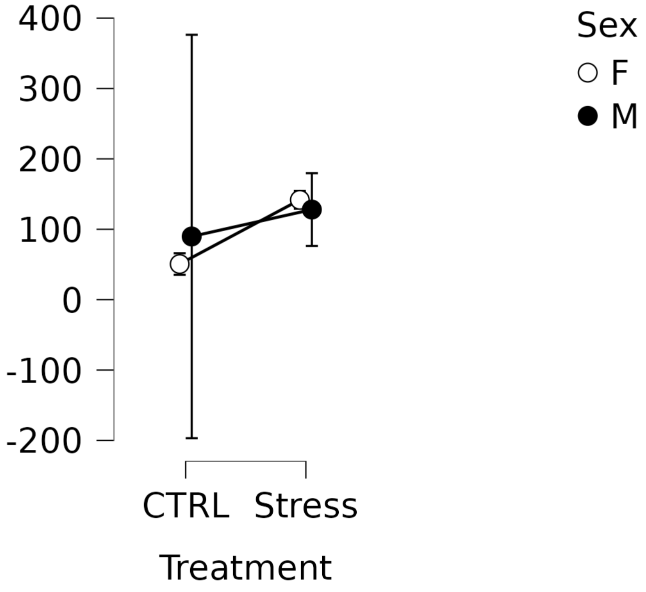


###### SPT Stage: SN

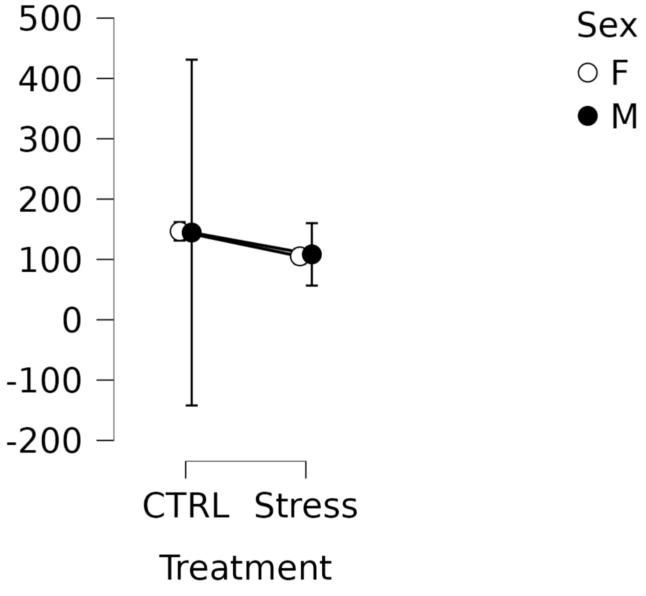

#### Post Hoc Tests

*Post Hoc Comparisons - Treatment*

| | | Mean Difference | SE | df | t | $p_{Holm}$ |
|---|---|---|---|---|---|---|
| CTRL | Stress | −12.96 | 9.311 | 33 | −1.391 | .173 |

*Note.* Results are averaged over the levels of: Sex, SPT Stage

*Letter-Based Grouping - Treatment*

| Treatment | Letter |
|---|---|
| CTRL | a |
| Stress | a |

*Note.* If two or more means share the same grouping symbol, then we cannot show them to be different, but we also did not show them to be the same.

*Post Hoc Comparisons - SPT Stage*

| | | Mean Difference | SE | df | t | $p_{Holm}$ |
|---|---|---|---|---|---|---|
| SI | SN | −23.61 | 9.046 | 33 | −2.610 | .014* |

* $p < .05$

*Note.* Results are averaged over the levels of: Treatment, Sex

*Letter-Based Grouping - SPT Stage*

| SPT Stage | Letter |
|---|---|
| SI | a |
| SN | b |

*Note.* If two or more means share the same grouping symbol, then we cannot show them to be different, but we also did not show them to be the same.

*Post Hoc Comparisons - Treatment ✻ SPT Stage*

| | | Mean Difference | SE | df | t | $p_{Holm}$ |
|---|---|---|---|---|---|---|
| CTRL, SI | Stress, SI | −64.73 | 13.82 | 33 | −4.685 | < .001*** |
| | CTRL, SN | −75.38 | 14.92 | 33 | −5.053 | < .001*** |
| | Stress, SN | −36.57 | 13.29 | 33 | −2.752 | .029* |
| Stress, SI | CTRL, SN | −10.66 | 12.67 | 33 | −0.8412 | .406 |
| | Stress, SN | 28.16 | 10.23 | 33 | 2.751 | .029* |
| CTRL, SN | Stress, SN | 38.81 | 12.09 | 33 | 3.210 | .012* |

* p < .05, *** p < .001

*Note.* P-value adjusted for comparing a family of 6 estimates.

*Note.* Results are averaged over the levels of: Sex

*Letter-Based Grouping - Treatment:SPT Stage*

| Treatment | SPT Stage | Letter |
|---|---|---|
| CTRL | SI | a |
| Stress | | c |
| CTRL | SN | c |
| Stress | | b |

*Note.* If two or more means share the same grouping symbol,
then we cannot show them to be different, but we also did not show them to be the same.

### RMANOVA - Erratic swimming

*Within Subjects Effects*

| Cases | Sum of Squares | df | Mean Square | F | p | $\eta^2_p$ |
|---|---|---|---|---|---|---|
| SPT Stage | 18.57 | 1 | 18.57 | 0.5972 | .445 | 0.01777 |
| SPT Stage ∗ Treatment | 14.05 | 1 | 14.05 | 0.4519 | .506 | 0.01351 |
| SPT Stage ∗ Sex | 0.8036 | 1 | 0.8036 | 0.02585 | .873 | $7.827 \times 10^{-4}$ |
| SPT Stage ∗ Treatment ∗ Sex | 0.5560 | 1 | 0.5560 | 0.01788 | .894 | $5.416 \times 10^{-4}$ |
| Residuals | 1026 | 33 | 31.09 | | | |

*Note.* Type III Sum of Squares

*Between Subjects Effects*

| Cases | Sum of Squares | df | Mean Square | F | p | $\eta^2_p$ |
|---|---|---|---|---|---|---|
| Treatment | 583.8 | 1 | 583.8 | 30.64 | < .001 | 0.4815 |
| Sex | 0.1979 | 1 | 0.1979 | 0.01039 | .919 | $3.147 \times 10^{-4}$ |
| Treatment ∗ Sex | 11.32 | 1 | 11.32 | 0.5944 | .446 | 0.01769 |
| Residuals | 628.7 | 33 | 19.05 | | | |

*Note.* Type III Sum of Squares

#### Descriptives

##### Descriptives plots

###### SPT Stage: SI

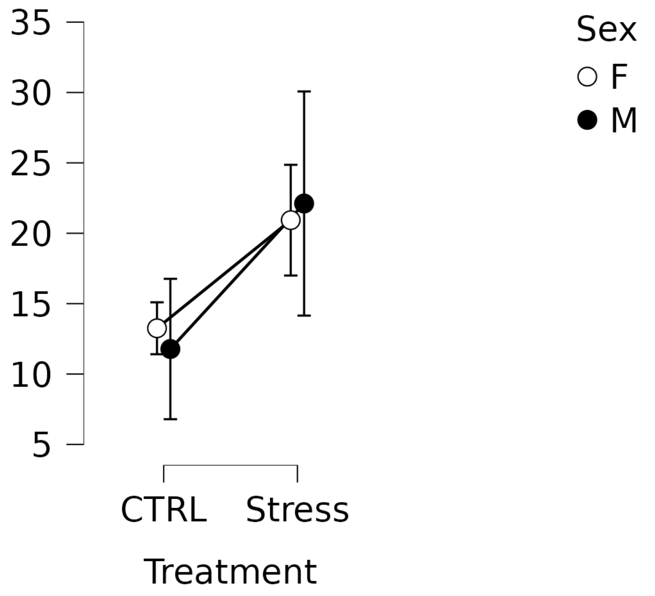


###### SPT Stage: SN

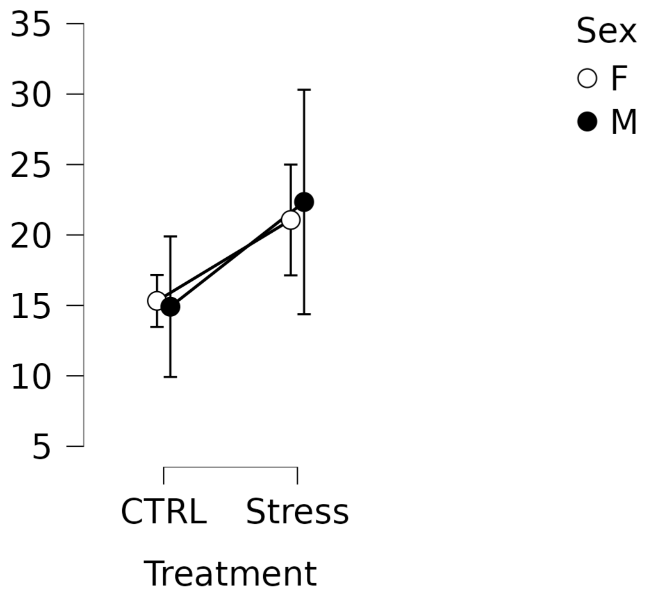

#### Post Hoc Tests

*Post Hoc Comparisons - Treatment*

| | | Mean Difference | SE | df | t | $p_{Holm}$ |
|---|---|---|---|---|---|---|
| CTRL | Stress | −7.798 | 1.409 | 33 | −5.535 | < .001*** |

*** p < .001

*Note.* Results are averaged over the levels of: Sex, SPT Stage

*Letter-Based Grouping - Treatment*

| Treatment | Letter |
|---|---|
| CTRL | a |
| Stress | b |

*Note.* If two or more means share the same grouping symbol, then we cannot show them to be different, but we also did not show them to be the same.

*Post Hoc Comparisons - SPT Stage*

| | | Mean Difference | SE | df | t | $p_{Holm}$ |
|---|---|---|---|---|---|---|
| SI | SN | −1.391 | 1.800 | 33 | −0.7728 | .445 |

*Note.* Results are averaged over the levels of: Treatment, Sex

*Letter-Based Grouping - SPT Stage*

| SPT Stage | Letter |
|---|---|
| SI | a |
| SN | a |

*Note.* If two or more means share the same grouping symbol, then we cannot show them to be different, but we also did not show them to be the same.

*Post Hoc Comparisons - Treatment ✻ SPT Stage*

| | | Mean Difference | SE | df | t | $p_{Holm}$ |
|---|---|---|---|---|---|---|
| CTRL, SI | Stress, SI | −9.008 | 2.229 | 33 | −4.041 | .002** |
| | CTRL, SN | −2.600 | 2.968 | 33 | −0.8761 | .775 |
| | Stress, SN | −9.189 | 2.265 | 33 | −4.056 | .002** |
| Stress, SI | CTRL, SN | 6.407 | 2.305 | 33 | 2.779 | .033* |
| | Stress, SN | −0.1809 | 2.036 | 33 | −0.08887 | .930 |
| CTRL, SN | Stress, SN | −6.588 | 2.340 | 33 | −2.815 | .033* |

* p < .05, ** p < .01

*Note.* P-value adjusted for comparing a family of 6 estimates.

*Note.* Results are averaged over the levels of: Sex

*Letter-Based Grouping - Treatment:SPT Stage*

| Treatment | SPT Stage | Letter |
|---|---|---|
| CTRL | SI | a |
| Stress | | b |
| CTRL | SN | a |
| Stress | | b |

*Note.* If two or more means share the same grouping symbol,
then we cannot show them to be different, but we also did not show them to be the same.

### RMANOVA - Distance to bottom

*Within Subjects Effects*

| Cases | Sum of Squares | df | Mean Square | F | p | $\eta^2_p$ |
|---|---|---|---|---|---|---|
| SPT Stage | 4.822 | 1 | 4.822 | 1.788 | .190 | 0.05140 |
| SPT Stage ∗ Treatment | 4.910 | 1 | 4.910 | 1.821 | .186 | 0.05229 |
| SPT Stage ∗ Sex | 4.777 | 1 | 4.777 | 1.772 | .192 | 0.05095 |
| SPT Stage ∗ Treatment ∗ Sex | 7.219 | 1 | 7.219 | 2.677 | .111 | 0.07504 |
| Residuals | 88.99 | 33 | 2.697 | | | |

*Note.* Type III Sum of Squares

*Between Subjects Effects*

| Cases | Sum of Squares | df | Mean Square | F | p | $\eta^2_p$ |
|---|---|---|---|---|---|---|
| Treatment | 104.1 | 1 | 104.1 | 22.92 | < .001 | 0.4098 |
| Sex | 7.886 | 1 | 7.886 | 1.737 | .197 | 0.05000 |
| Treatment ∗ Sex | 7.562 | 1 | 7.562 | 1.665 | .206 | 0.04804 |
| Residuals | 149.8 | 33 | 4.541 | | | |

*Note.* Type III Sum of Squares

#### Descriptives

##### Descriptives plots

###### SPT Stage: SI

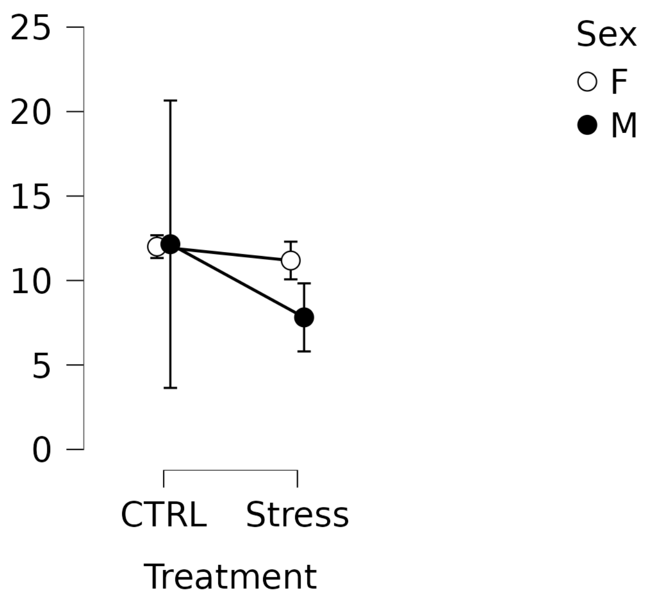


###### SPT Stage: SN

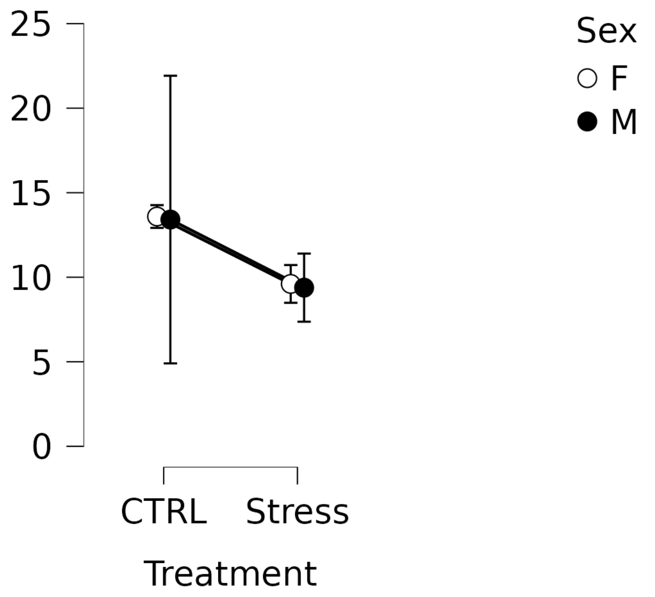

#### Post Hoc Tests

*Post Hoc Comparisons - Treatment*

| | | Mean Difference | SE | df | t | $p_{Holm}$ |
|---|---|---|---|---|---|---|
| CTRL | Stress | 3.292 | 0.6877 | 33 | 4.787 | < .001*** |

*** p < .001

*Note.* Results are averaged over the levels of: Sex, SPT Stage

*Letter-Based Grouping - Treatment*

| Treatment | Letter |
|---|---|
| CTRL | b |
| Stress | a |

*Note.* If two or more means share the same grouping symbol, then we cannot show them to be different, but we also did not show them to be the same.

*Post Hoc Comparisons - SPT Stage*

| | | Mean Difference | SE | df | t | $p_{Holm}$ |
|---|---|---|---|---|---|---|
| SI | SN | −0.7087 | 0.5300 | 33 | −1.337 | .190 |

*Note.* Results are averaged over the levels of: Treatment, Sex

*Letter-Based Grouping - SPT Stage*

| SPT Stage | Letter |
|---|---|
| SI | a |
| SN | a |

*Note.* If two or more means share the same grouping symbol, then we cannot show them to be different, but we also did not show them to be the same.

*Post Hoc Comparisons - Treatment ✻ SPT Stage*

| | | Mean Difference | SE | df | t | $p_{Holm}$ |
|---|---|---|---|---|---|---|
| CTRL, SI | Stress, SI | 2.577 | 0.6696 | 33 | 3.849 | .002** |
| | CTRL, SN | −1.424 | 0.8741 | 33 | −1.629 | .226 |
| | Stress, SN | 2.584 | 0.8024 | 33 | 3.220 | .009** |
| Stress, SI | CTRL, SN | −4.001 | 0.9294 | 33 | −4.305 | < .001*** |
| | Stress, SN | 0.006407 | 0.5996 | 33 | 0.01069 | .992 |
| CTRL, SN | Stress, SN | 4.008 | 1.029 | 33 | 3.894 | .002** |

* p < .05, ** p < .01, *** p < .001

*Note.* P-value adjusted for comparing a family of 6 estimates.

*Note.* Results are averaged over the levels of: Sex

*Letter-Based Grouping - Treatment:SPT Stage*

| Treatment | SPT Stage | Letter |
|---|---|---|
| CTRL | SI | b |
| Stress | | a |
| CTRL | SN | b |
| Stress | | a |

*Note.* If two or more means share the same grouping symbol,
then we cannot show them to be different, but we also did not show them to be the same.

### RMANOVA - Swimming speed

*Within Subjects Effects*

| Cases | Sum of Squares | df | Mean Square | F | p | $\eta^2_p$ |
|---|---|---|---|---|---|---|
| SPT Stage | 2.953 | 1 | 2.953 | 0.4576 | .503 | 0.01368 |
| SPT Stage ∗ Treatment | 4.048 | 1 | 4.048 | 0.6273 | .434 | 0.01865 |
| SPT Stage ∗ Sex | 0.6871 | 1 | 0.6871 | 0.1065 | .746 | 0.003216 |
| SPT Stage ∗ Treatment ∗ Sex | 2.209 | 1 | 2.209 | 0.3423 | .562 | 0.01027 |
| Residuals | 212.9 | 33 | 6.453 | | | |

*Note.* Type III Sum of Squares

*Between Subjects Effects*

| Cases | Sum of Squares | df | Mean Square | F | p | $\eta^2_p$ |
|---|---|---|---|---|---|---|
| Treatment | 87.12 | 1 | 87.12 | 11.70 | .002 | 0.2617 |
| Sex | 12.59 | 1 | 12.59 | 1.690 | .203 | 0.04873 |
| Treatment ∗ Sex | 0.8950 | 1 | 0.8950 | 0.1202 | .731 | 0.003628 |
| Residuals | 245.8 | 33 | 7.448 | | | |

*Note.* Type III Sum of Squares

#### Descriptives

##### Descriptives plots

###### SPT Stage: SI

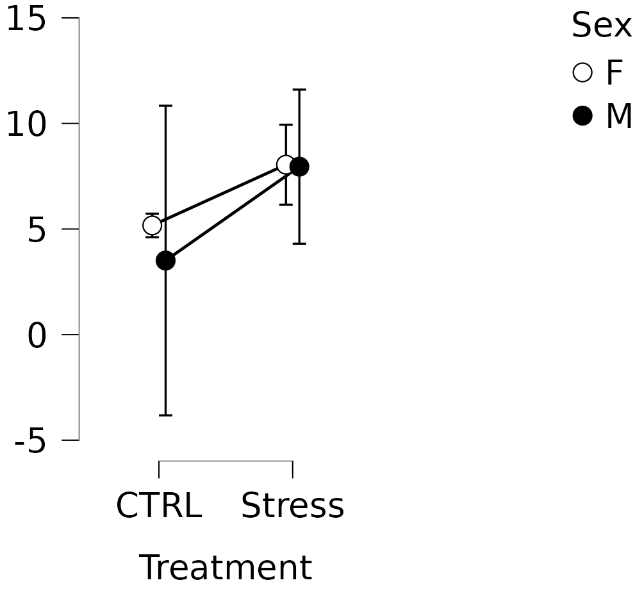


###### SPT Stage: SN

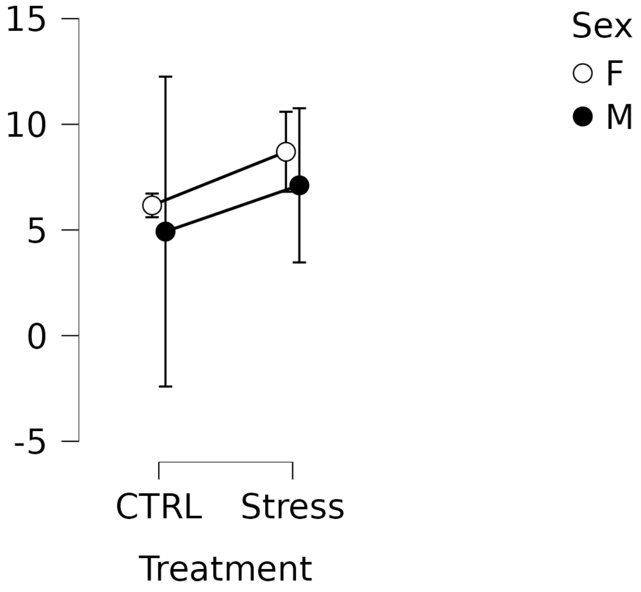

#### Post Hoc Tests

*Post Hoc Comparisons - Treatment*

| | | Mean Difference | SE | df | t | $p_{Holm}$ |
|---|---|---|---|---|---|---|
| CTRL | Stress | −3.012 | 0.8808 | 33 | −3.420 | .002** |

** $p < .01$

*Note.* Results are averaged over the levels of: Sex, SPT Stage

*Letter-Based Grouping - Treatment*

| Treatment | Letter |
|---|---|
| CTRL | a |
| Stress | b |

*Note.* If two or more means share the same grouping symbol, then we cannot show them to be different, but we also did not show them to be the same.

*Post Hoc Comparisons - SPT Stage*

| | | Mean Difference | SE | df | t | $p_{Holm}$ |
|---|---|---|---|---|---|---|
| SI | SN | −0.5546 | 0.8199 | 33 | −0.6765 | .503 |

*Note.* Results are averaged over the levels of: Treatment, Sex

*Letter-Based Grouping - SPT Stage*

| SPT Stage | Letter |
|---|---|
| SI | a |
| SN | a |

*Note.* If two or more means share the same grouping symbol, then we cannot show them to be different, but we also did not show them to be the same.

*Post Hoc Comparisons - Treatment ✻ SPT Stage*

| | | Mean Difference | SE | df | t | $p_{Holm}$ |
|---|---|---|---|---|---|---|
| CTRL, SI | Stress, SI | −3.662 | 1.220 | 33 | −3.001 | .031* |
| | CTRL, SN | −1.204 | 1.352 | 33 | −0.8904 | .759 |
| | Stress, SN | −3.567 | 1.209 | 33 | −2.949 | .031* |
| Stress, SI | CTRL, SN | 2.458 | 1.197 | 33 | 2.053 | .192 |
| | Stress, SN | 0.09473 | 0.9276 | 33 | 0.1021 | .919 |
| CTRL, SN | Stress, SN | −2.363 | 1.186 | 33 | −1.992 | .192 |

* $p < .05$

*Note.* P-value adjusted for comparing a family of 6 estimates.

*Note.* Results are averaged over the levels of: Sex

*Letter-Based Grouping - Treatment:SPT Stage*

| Treatment | SPT Stage | Letter |
|---|---|---|
| CTRL | SI | a |
| Stress | | b |
| CTRL | SN | ab |
| Stress | | b |

*Note.* If two or more means share the same grouping symbol, then we cannot show them to be different, but we also did not show them to be the same.
